# Absolute quantitation of individual SARS-CoV-2 RNA molecules: a new paradigm for infection dynamics and variant differences

**DOI:** 10.1101/2021.06.29.450133

**Authors:** Jeffrey Y Lee, Peter AC Wing, Dalia S Gala, Marko Noerenberg, Aino I Järvelin, Josh Titlow, Xiaodong Zhuang, Natasha Palmalux, Louisa Iselin, Mary Kay Thompson, Richard M Parton, Alan Wainman, Daniel Agranoff, William James, Alfredo Castello, Jane A McKeating, Ilan Davis

## Abstract

Despite an unprecedented global research effort on SARS-CoV-2, early replication events remain poorly understood. Given the clinical importance of emergent viral variants with increased transmission, there is an urgent need to understand the early stages of viral replication and transcription. We used single molecule fluorescence *in situ* hybridisation (smFISH) to quantify positive sense RNA genomes with 95% detection efficiency, while simultaneously visualising negative sense genomes, sub-genomic RNAs and viral proteins. Our absolute quantification of viral RNAs and replication factories revealed that SARS-CoV-2 genomic RNA is long-lived after entry, suggesting that it avoids degradation by cellular nucleases. Moreover, we observed that SARS-CoV-2 replication is highly variable between cells, with only a small cell population displaying high burden of viral RNA. Unexpectedly, the B.1.1.7 variant, first identified in the UK, exhibits significantly slower replication kinetics than the Victoria strain, suggesting a novel mechanism contributing to its higher transmissibility with important clinical implications.

**Graphical Abstract:** 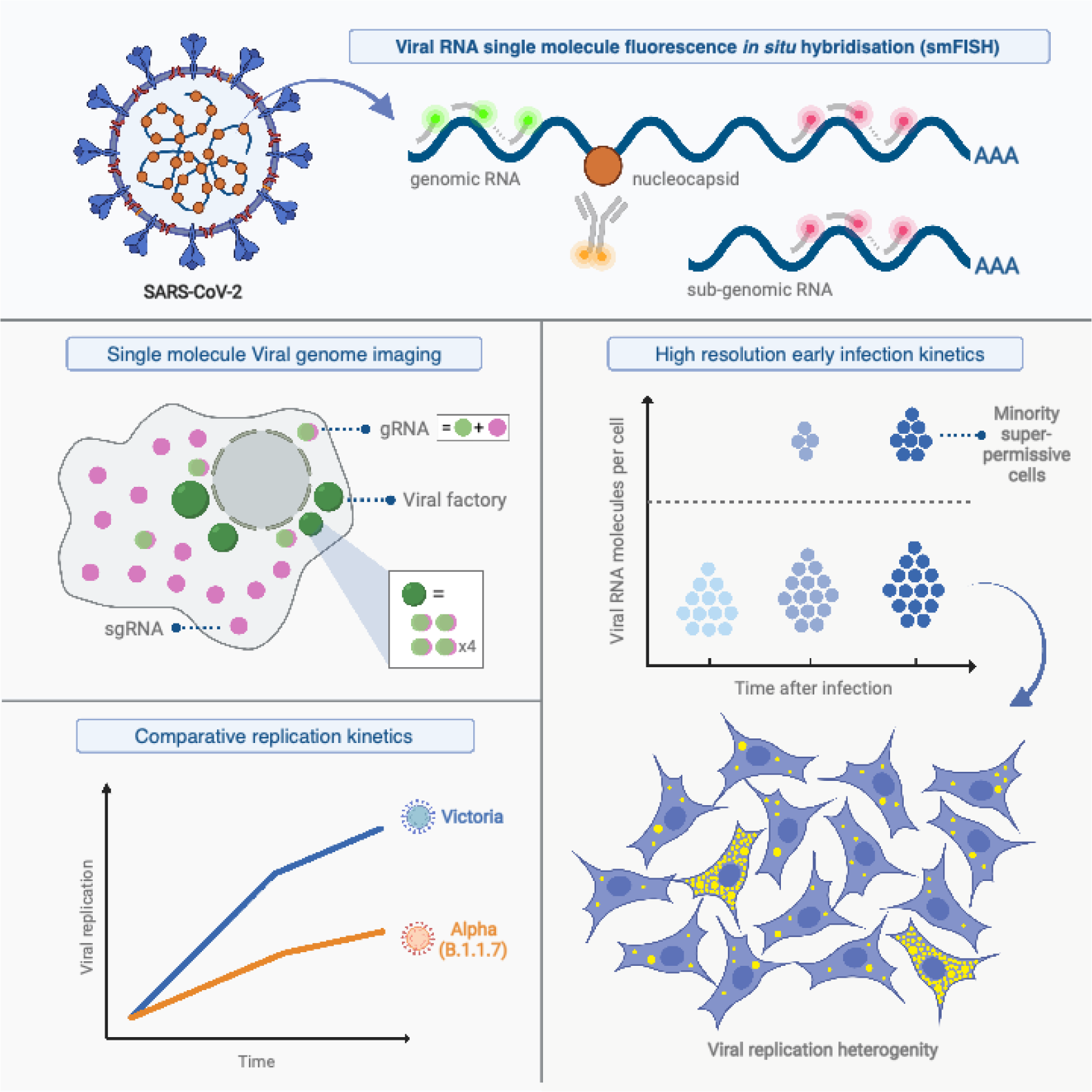

**In brief:** By detecting nearly all individual SARS-CoV-2 RNA molecules, we quantified viral replication and defined cell susceptibility to infection. We discovered that a minority of cells show significantly elevated viral RNA levels and observed slower replication kinetics for the Alpha variant relative to the Victoria strain.

**Highlights:**

- Single molecule quantification of SARS-CoV-2 replication uncovers early infection kinetics
- There is substantial heterogeneity between cells in rates of SARS-CoV-2 replication
- Genomic RNA is stable and persistent during the initial stages of infection
- B.1.1.7 variant replicates more slowly than the Victoria strain

## Introduction

Severe acute respiratory syndrome coronavirus 2 (SARS-CoV-2) is the causative agent of the COVID-19 pandemic. The viral genome consists of a single positive strand genomic RNA (+gRNA) approximately 30kb in length that encodes a plethora of viral proteins (Kim et al., 2020; Zhao et al., 2021). SARS-CoV-2 primarily targets the respiratory tract and infection is mediated by Spike protein binding to human angiotensin-converting enzyme (ACE2), where the transmembrane protease serine 2 (TMPRSS2) triggers fusion of the viral and cell membranes (Hoffmann et al., 2020; Wan et al., 2020). Following virus entry and capsid trafficking to the endoplasmic reticulum, the first step in the replicative life cycle is the translation of the gRNA to synthesise the replicase complex. This complex synthesises the negative sense genomic strand, enabling the production of additional positive gRNA copies. In addition, a series of shorter sub-genomic RNAs (sgRNAs) are synthesised that encode the structural matrix, spike, nucleocapsid and envelope proteins as well as a series of non-structural proteins (Kim *et al*., 2020; Sola et al., 2015). The intracellular localisation of these early events were described using electron microscopy (Laue et al., 2021) and by antibody-based imaging of viral double stranded (ds)RNA (Lean et al., 2020). However, the J2 dsRNA antibody lacks sensitivity and specificity at early times post infection, as the low abundance of SARS-CoV-2 dsRNA is indistinguishable from host dsRNAs (Dhir et al., 2018). Our current knowledge of these early steps in the SARS-CoV-2 replicative life cycle are poorly understood despite their essential role in the establishment of productive infection.

Since the initial outbreak in the Wuhan province of China in 2019 several geographically distinct variants of concern (VOC) with altered transmission have arisen (Chen et al., 2020; Lythgoe et al., 2021). Emerging VOC such as the recently named Alpha strain (previously known and referred to herein as B.1.1.7), first detected in Kent in the UK, possess a fitness advantage in terms of their ability to transmit compared to the Victoria (VIC) isolate, an early strain of SARS-CoV-2 first detected in Wuhan in China (Caly et al., 2020; Davies et al., 2021; Kidd et al., 2021; Volz et al., 2021). Many of the VOC encode mutations in the Spike (S) protein (Rees-Spear et al., 2021) and, consequently, the effects of these amino acid substitutions on viral entry and immuno-evasion are under intense study (Kissler et al., 2021; Washington et al., 2021). However, some of the mutations map to non-structural proteins, so could impact viral replication dynamics. To date, the early replication events of SARS-CoV-2 variants have not been characterised, as the current techniques for quantifying SARS-CoV-2 genomes and replication rates rely on bulk approaches or have limited sensitivity.

The use of single molecule and single-cell analyses in biology offers unprecedented insights into the behaviour of individual cells and the stochastic nature of gene expression that are often masked by population-based studies (Fraser and Kaern, 2009; Raj and van Oudenaarden, 2009). These approaches have revealed how cells vary in their ability to support viral growth and how stochastic forces can inform our understanding of the infection process (Billman et al., 2017; Boersma et al., 2020; Chou and Lionnet, 2018; Shulla and Randall, 2015; Singer et al., 2021). Fluorescence in situ hybridisation (FISH) was previously used to detect RNAs in hepatitis C virus and Sindbis infected cells with high sensitivity (Garcia-Moreno et al., 2019; Ramanan et al., 2016; Singer *et al*., 2021). This approach has been applied to SARS-CoV-2 in a limited capacity (Burke et al., 2021; Rensen et al., 2021) with most studies utilising amplification-based signal detection methods to visualise viral RNA (Best Rocha et al., 2020; Carossino et al., 2020; Guerini-Rocco et al., 2020; Jiao et al., 2020; Kusmartseva et al., 2020; Lean *et al*., 2020; Liu et al., 2020). These experiments used either chromogenic histochemical detection using bright field microscopy, or detection of fluorescent dyes, which both lack the sensitivity to detect individual RNA molecules. Consequently, the kinetics of SARS-CoV-2 RNA replication and transcription during the early phase of infection are not well understood and lack quantitative, spatial and temporal information on the genesis of gRNA and sgRNAs. To address this gap, we developed a single molecule (sm)FISH method based on earlier published protocols (Femino et al., 1998; Raj et al., 2008; Singer *et al*., 2021; Titlow et al., 2018) to visualise SARS-CoV-2 RNAs with high sensitivity and spatial precision, providing a powerful new approach to track infection through the detection and quantification of viral replication factories. Our results uncover a previously unrecognised heterogeneity among cells in supporting SARS-CoV-2 replication and a surprisingly slower replication rate of the B.1.1.7 variant when compared to the early lineage VIC strain.

## Results

### SARS-CoV-2 genomic RNA at single molecule resolution

To explore the spatial and temporal aspects of SARS-CoV-2 replication at single molecule and cell levels, we carried out smFISH experiments with fluorescently labelled probes directed against the 30kb viral gRNA. 48 short antisense DNA oligonucleotide probes were designed to target the viral ORF1a and labelled with a single fluorescent dye to detect the positive sense gRNA, as described previously (Gaspar et al., 2017) (Figure 1A). The probe set detected single molecules of gRNA within SARS-CoV-2 infected Vero E6 cells, visible as well-resolved diffraction-limited single spots with a consistent fluorescence intensity and shape (Figure 1B). Treatment of the infected cells with RNase or the viral polymerase inhibitor remdesivir (RDV) ablated the probe signal, confirming specificity (Figure S1A). To assess the efficiency and specificity of detection of the +ORF1a probe set, we divided the probes into two groups of 24 alternating oligonucleotides (“ODD” and “EVEN”) that were labelled with different fluorochromes. Interlacing the probes minimised chromatic aberration between spots detected by the two colours (Figure 1C). Analysis of the SARS-CoV-2 gRNA with these probes showed a mean distance of <250nm between the two fluorescent spots, indicating near-perfect colour registration and a lack of chromatic aberration. 95% of the diffraction-limited spots within infected cells were dual labelled, demonstrating efficient detection of single SARS-CoV-2 gRNA molecules (Figure 1C).

**Figure 1.**
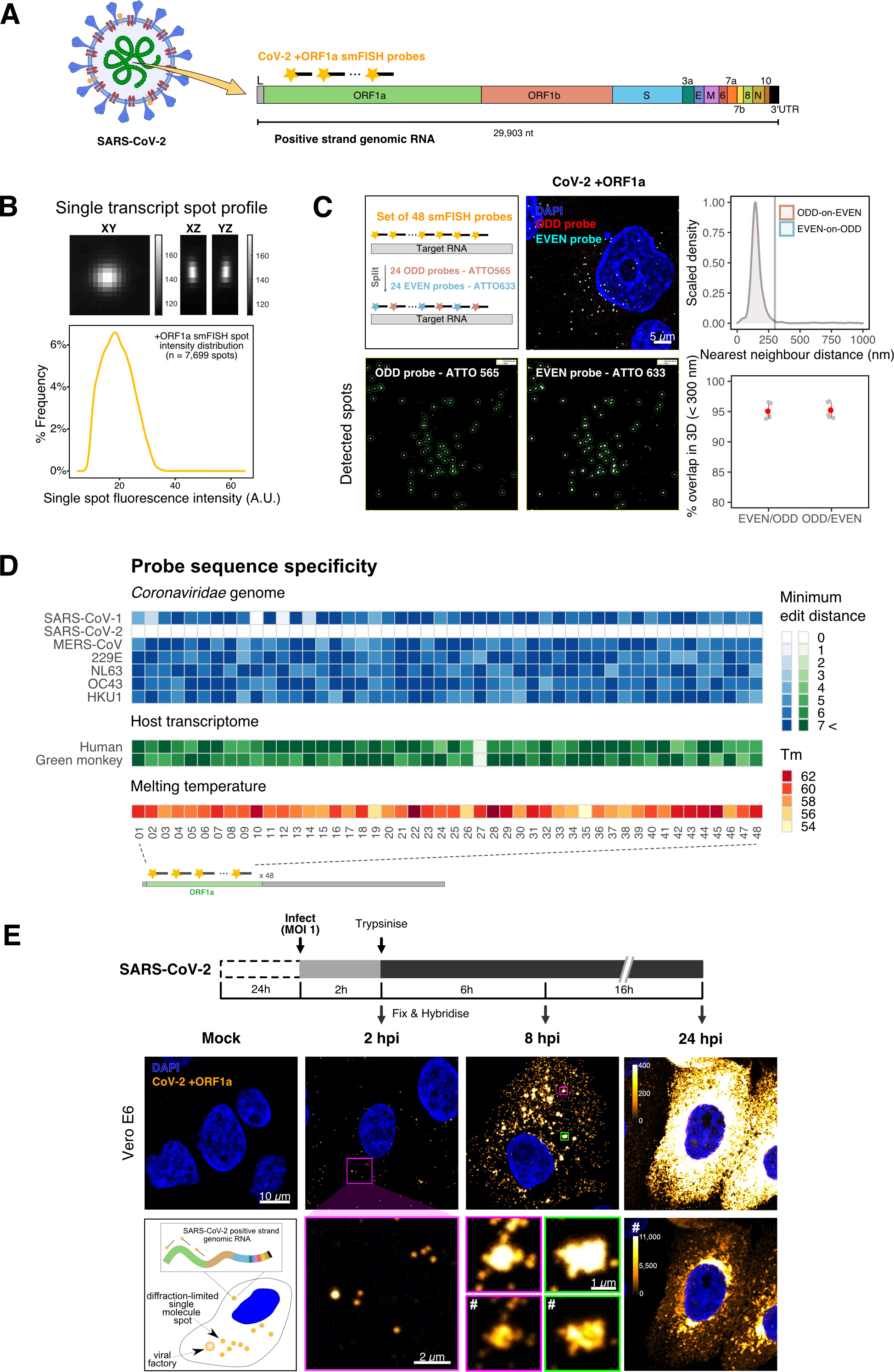
Sensitive single-molecule detection of SARS-CoV-2 genomic RNA in infected cells. (A) Schematic illustration of single-molecule fluorescence in situ hybridisation (smFISH) for detecting SARS-CoV-2 positive strand genomic RNA (+gRNA) within infected cells. (B) Reference spatial profile of a diffraction-limited +ORF1a smFISH spot. The calibration bar represents relative fluorescence intensity (top). Frequency distribution of smFISH spot intensities, exhibiting a unimodal distribution (bottom). (C) Assessment of smFISH detection sensitivity by a dual-colour co-detection method. Maximum intensity projected images and corresponding FISH-quant spot detection views of ODD and EVEN probe sets are shown. Scale bar = 5 µm. Density histogram of nearest-neighbour distance from one spectral channel to another (top). Vertical line indicates 300 nm distance. Percentage overlap between spots detected by ODD and EVEN split probes, calculated bidirectionally (bottom). (D) Heatmap of probe sequence alignment against various *Coronaviridae* and host transcriptomes. Each column represents individual 20 nt +ORF1a probe sequences. The minimum edit distance represents mismatch scores, where ‘0’ indicates a perfect match. Melting temperatures of each probe at the smFISH hybridisation condition are shown. (E) Experimental design for visualising SARS-CoV-2 gRNA with smFISH at different time points after infection of Vero E6 cells. Cells were seeded on cover-glass and 24 h later, inoculated with SARS-CoV-2 (VIC strain at MOI 1) for 2 h. Non-internalised viruses were removed by trypsin digestion and cells fixed at the timepoints shown. Representative 4 µm maximum intensity projection confocal images are shown. Images are contrasted according to the calibration bar in the upper panel of 24 hpi unless indicated with a symbol ‘#’ for wider dynamic contrast range. Magnified view of insets in the upper panels are shown in lower panels. Scale bars = 10 µm or 2 µm.

To verify the specificity of the +ORF1a probes for SARS-CoV-2, we aligned them against other coronaviruses and the human transcriptome. Many of the oligonucleotides showed mismatches with SARS-CoV-1, MERS and other coronaviruses along with human RNAs (Figure 1D). We also evaluated their ability to bind HCoV-229E RNA within infected Huh-7 cells. Although the J2 antibody, specific for double stranded (ds)RNA, detected dsRNA foci in the HCoV-229E infected cells, no signal was seen with the SARS-CoV-2 +ORF1a probe set (Figure S1B). Taken together, these results highlight the efficiency and specificity of the +ORF1a probe set for detecting single molecules of SARS-CoV-2 gRNA.

Having established smFISH for the detection of SARS-CoV-2 gRNA, we used this technique to assess both the quantity and distribution of gRNA during infection. Cells were inoculated with virus at a multiplicity of infection (MOI) of 1 for 2h and non-internalised virus removed by trypsin digestion to synchronise the infection. At 2 hours post infection (hpi), most fluorescent spots correspond to single gRNAs along with a small number of foci harbouring several gRNA copies (Figure 1E), consistent with early RNA replication events. By 8 hpi, we noted an expansion in the number of bright multi-gRNA foci and at 24 hpi there was a further increase in the number of multi-RNA foci that localised to the perinuclear region (Figure 1E); consistent with the reported association of viral replication factories with membranous structures derived from the endoplasmic reticulum (V’Kovski et al., 2021). Interestingly, our observation of individual gRNA molecules at the periphery of cells (Figure 1E) are also consistent with individual viral particles observed at the same location by electron microscopy (Cortese et al., 2020; Klein et al., 2020).

To assess whether virion-encapsulated RNA is accessible to the probes, we immobilized SARS-CoV-2 particles from our viral stocks on glass and incubated them with the +ORF1a probes. We observed a large number of spots in the immobilised virus preparation that was compatible with single RNA molecules (Figure S1C), suggesting that detection of RNA within viral particles was achieved. We conclude that detection of SARS-CoV-2 +gRNA by smFISH identifies changes in viral RNA abundance and cellular distribution during early replication that includes all the known locations of +gRNA molecules in the cell, namely in virions, free in the cytoplasm and within ER-like membranous structures.

### Quantification of SARS-CoV-2 genomic and sub-genomic RNAs

SARS-CoV-2 produces both gRNA and subgenomic (sg)RNAs which are both critical to its replicative life cycle. However, quantitation of sgRNAs is challenging due to their sequence overlap with the 3’ end of the gRNA. To estimate the abundance of sgRNAs we designed two additional probe sets labelled with different fluorochromes; an +ORF-N set that hybridises to all canonical positive sense viral RNAs, and an +ORF-S set that detects only sgRNA encoding S (S-sgRNA) and gRNA (Figure 2A) (Kim *et al*., 2020). Therefore, spots showing fluorescence only for +ORF-N or +ORF-S probe sets will represent sgRNAs, whereas spots positive for both +ORF-N or +ORF-S and +ORF1a will correspond to gRNA molecules. We applied this approach to visualise SARS-CoV-2 RNAs in infected Vero E6 cells (6 hpi) and observed a high abundance of sgRNAs compared to gRNAs (Figure 2B), in agreement with RNA sequencing studies (Alexandersen et al., 2020; Kim *et al*., 2020). Further analysis revealed that the +ORF-N and +ORF-S single labelled spots, corresponding to sgRNAs, were more uniformly distributed throughout the cytoplasm than dual labelled gRNA, consistent with their predominant role as mRNAs to direct protein synthesis (Figure 2B). Association of gRNA with nucleocapsid (N) is essential for the assembly of coronavirus particles (Carlson et al., 2020; Dinesh et al., 2020; Iserman et al., 2020). To monitor this process in SARS-CoV-2, we combined smFISH using the +ORF1a and +ORF-N probe sets with immunofluorescence detection of the viral nucleocapsid (N). Our findings show that N protein primarily co-localises with gRNA, while displaying a limited overlap with sgRNAs (Figure 2C). Together, these data demonstrate the specificity of our probes to accurately discriminate between the viral genomic and sub-genomic RNA forms.

**Figure 2.**
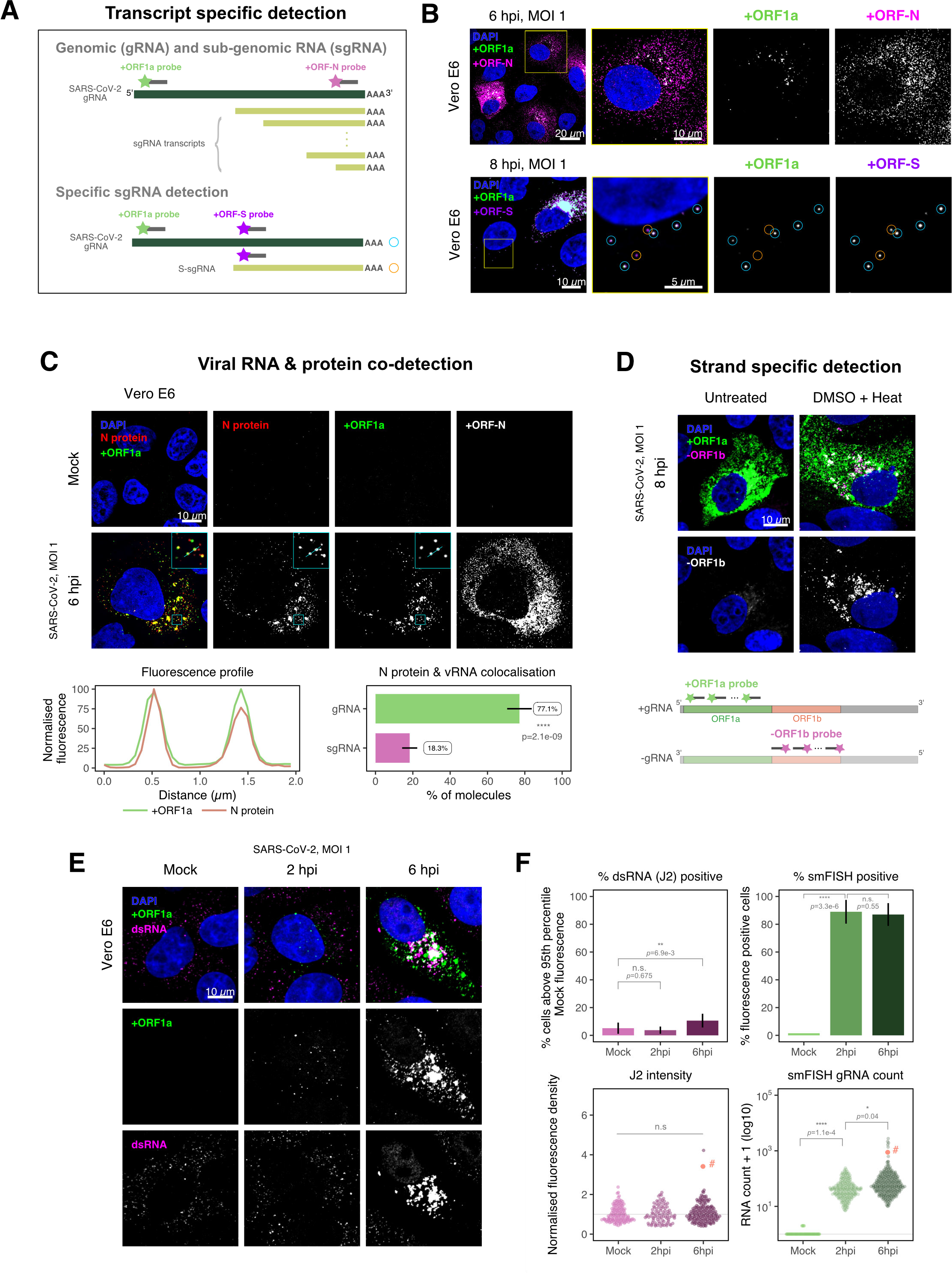
Dissecting SARS-CoV-2 gene expression using smFISH. (A) Schematic illustration of transcript specific targeting of SARS-CoV-2 genomic RNA (gRNA) and subgenomic RNA (sgRNA) using smFISH. (B) Transcript specific visualisation of gRNA and sgRNA in infected (VIC strain) Vero E6 cells. Cells were infected as in (Figure 1D) and hybridised with probes against +ORF1a & +ORF-N probe at 6 hpi (upper panels) or +ORF1a & +ORF-S probe at 8 hpi (lower panels). Representative 3 µm maximum intensity projected confocal images are shown. Orange circular regions of interest (ROIs) indicate S-sgRNA encoding Spike whereas dual-colour spots (teal-coloured ROIs) represent gRNA. Scale bar = 5, 10 or 20 µm. (C) Co-detection of viral nucleocapsid (N) with gRNA and sgRNA. Monoclonal anti-N (Ey2B clone) was used for N protein immunofluorescence. Representative 3 µm z-projected confocal images are shown. The inset shows a magnified view of co-localised N and gRNA. Scale bar = 10 µm. Fluorescence profiles of N immunostaining and gRNA smFISH intensity across a 2 µm linear distance are shown in the image inset (lower left). Percentage of co-localised gRNA or sgRNA molecules with N-protein at 6 hpi. Co-localisation was assessed by N fluorescence density within point-spread function ellipsoids of RNA spots over random coordinates. sgRNA were defined as single-coloured spots with +ORF-N probe signal only (n=7) (lower right). Student’s t-test. ****, p<0.0001. (D) Detection of both positive and negative genomic RNA by denaturing viral dsRNA with DMSO and heat treatment at 80°C (upper panels). 3 µm z-projected images of infected Vero E6 cells at 8 hpi are shown. Scale bar = 10 µm. Schematic illustration of +ORF1a and -ORF1b probe targeting regions (lower panel). -ORF1b probe target region does not overlap with +ORF1a target sequences to prevent probe duplex formation. (E) Comparison of anti-dsRNA (J2) and gRNA smFISH. Full z-projected images of infected Vero E6 cells co-stained with J2 and smFISH are shown. Scale bar = 10 µm. (F) Percentage of infected cells detected by J2 or smFISH (upper panels). For J2-based quantification, we defined the threshold as 95th percentile fluorescent signal of uninfected cells (Mock) due to the presence of endogenous host-derived signals. Fluorescent positive signals were used for smFISH-based quantification. Data are presented as mean ± S.D. Comparison of quantification results between J2 stain and smFISH (lower panels). Each symbol represents one cell. J2 signal was quantified by fluorescence density over 3D cell volume, which was normalised to the average signal of uninfected control cells (horizontal dotted line). gRNA count represents sum of single-molecule spots and decomposed spots within viral factories. The symbol denoted with ‘#’ is the infected cell shown in Figure 2E. (J2 stain, n=3 independent repeats; smFISH, n=4). One-way ANOVA and Tukey post-hoc test. n.s, not significant; *, p<0.05; **, p<0.01; ****, p<0.0001.

Negative sense gRNA and sgRNAs are the templates for the synthesis of positive sense RNAs and are expected to localise to viral replication factories. However, their detection by RT-qPCR or sequencing is hampered by cDNA library protocols that employ oligo(dT) selection and by primer binding to double stranded RNA (dsRNA) structures (Ramanan *et al*., 2016; Sethna et al., 1991). To detect negative sense viral RNAs, we denatured dsRNA complexes through either formamide, DMSO or sodium hydroxide treatment (Singer *et al*., 2021; Wilcox et al., 2019). The combination of DMSO with heat treatment resulted in a loss of anti-dsRNA J2 signal, while maintaining cell integrity, suggesting a disruption of dsRNA hybrids (Figure S2A). We designed an smFISH probe set specific for the ORF1b antisense sequence that targets the negative sense gRNA (-gRNA) and resulted in intense diffraction-limited spots in DMSO and heat-treated cells (Figure 2D). The -gRNA spots were detected at a significantly lower level than their +gRNA counterparts, with substantial overlap observed between the two strands at multi-RNA spots, consistent with these foci representing active sites of viral replication. To determine if these multi-RNA foci contain dsRNA, the permeabilised infected cells were treated with RNaseT1 or RNaseIII, which are nucleases specific for single-stranded RNA (ssRNA) and dsRNA, respectively (Figure S2B). RNaseT1 digestion diminished the +ORF1a probe signal, while RNaseIII treatment abolished the anti-dsRNA J2 signal. A cocktail of RNaseT1 and RNaseIII ablated both +ORF1a probe binding and anti-dsRNA J2 signals, demonstrating that the +ORF1a probe set hybridises to both single and duplex RNA under our experimental conditions (Figure S2B). Furthermore, treating cells with DMSO prior to RNaseT1 fully ablated the smFISH signal (Figure S2C), demonstrating that denaturation makes dsRNA accessible for RNaseT1 degradation. In summary, our data show that probe binding to negative strand gRNA requires chemical denaturation, suggesting that this replication intermediate localise within dsRNA structures.

### Anti-dsRNA antibodies underestimate SARS-CoV-2 replication

The establishment of replication factories is a critical phase of the virus life cycle. Previous reports have identified these viral factories using the J2 dsRNA antibody (Burgess and Mohr, 2015; Cortese *et al*., 2020; Targett-Adams et al., 2008; Weber et al., 2006). However, this approach depends on high levels of viral dsRNA as cells naturally express endogenous low levels of dsRNA (Dhir *et al*., 2018; Kimura et al., 2018) (Figure 2E). To evaluate the ability of J2 antibody to quantify SARS-CoV-2 replication sites, we co-stained infected cells at 2 and 6 hpi with both J2 and +ORF1a smFISH probes. No viral-specific J2 signal was detected at 2 hpi and only 10% of infected cells stained positive at 6 hpi, in agreement with previous observations (Cortese *et al*., 2020; Eymieux et al., 2021) (Figure 2E). In contrast, more than 85% of the cells showed diffraction-limited smFISH signals at both time points (Figure 2F). Furthermore, the average J2 signal detected in the SARS-CoV-2 infected cells at both time points was comparable to uninfected cells (Figure 2F). These data clearly show the J2 antibody, although broadly used, underestimates the frequency of SARS-CoV-2 infection. In contrast, smFISH detected gRNA as early as 2 hpi, with a significant increase in copy number by 6 hpi, highlighting its utility to detect and quantify viral replication factories.

### SARS-CoV-2 replication at single molecule resolution

The efficiency and sensitivity of the smFISH approach to detect single molecules of SARS-CoV-2 RNA allowed us to investigate the dynamics of viral replication in Vero E6 cells during the first 10h of infection (Figure 3A). At 2 hpi the +ORF1a probe set detected predominantly single molecules of +gRNA with a median value of ∼30 molecules per cell (Figure 3B-C). Of note, gRNA copies/cell at 2hpi were unaffected by RDV treatment, suggesting that these RNAs derive from incoming viral particles (Figure 3C). In contrast, the increase in gRNA copies/cell at 4 and 6 hpi was inhibited by RDV, reflecting active viral replication. The infected cell population showed varying gRNA levels that we classified into three groups; (i) ‘partially resistant’ cells with <10^2^ gRNA copies that showed no increase between 2 and 8 hpi (60% of the population); (ii) ‘permissive’ cells with ∼10^2^-10^5^ copies/cell showing a modest increase in gRNA burden over time (∼30%) and (iii) ‘super-permissive’ cells with >10^5^ copies/cell showing a dramatic increase in gRNA copies (∼10%). RDV treatment delayed the appearance of super-permissive cells until late times post infection (10-24 hpi) (Figure 3C-E). RDV inhibition of viral replication was partial despite using a concentration that exceeded the IC_90_ (Figure S3A). Analysing the total cellular gRNA content showed the dominant source of vRNA to be the ‘super-permissive’ cells (Figure 3D), suggesting that bulk RNA analyses such as RT-qPCR are biased by these group of cells containing high levels of gRNA. In the presence of RDV the average gRNA copies/cell were stable over time, suggesting that incoming gRNA is long lived.

**Figure 3.**
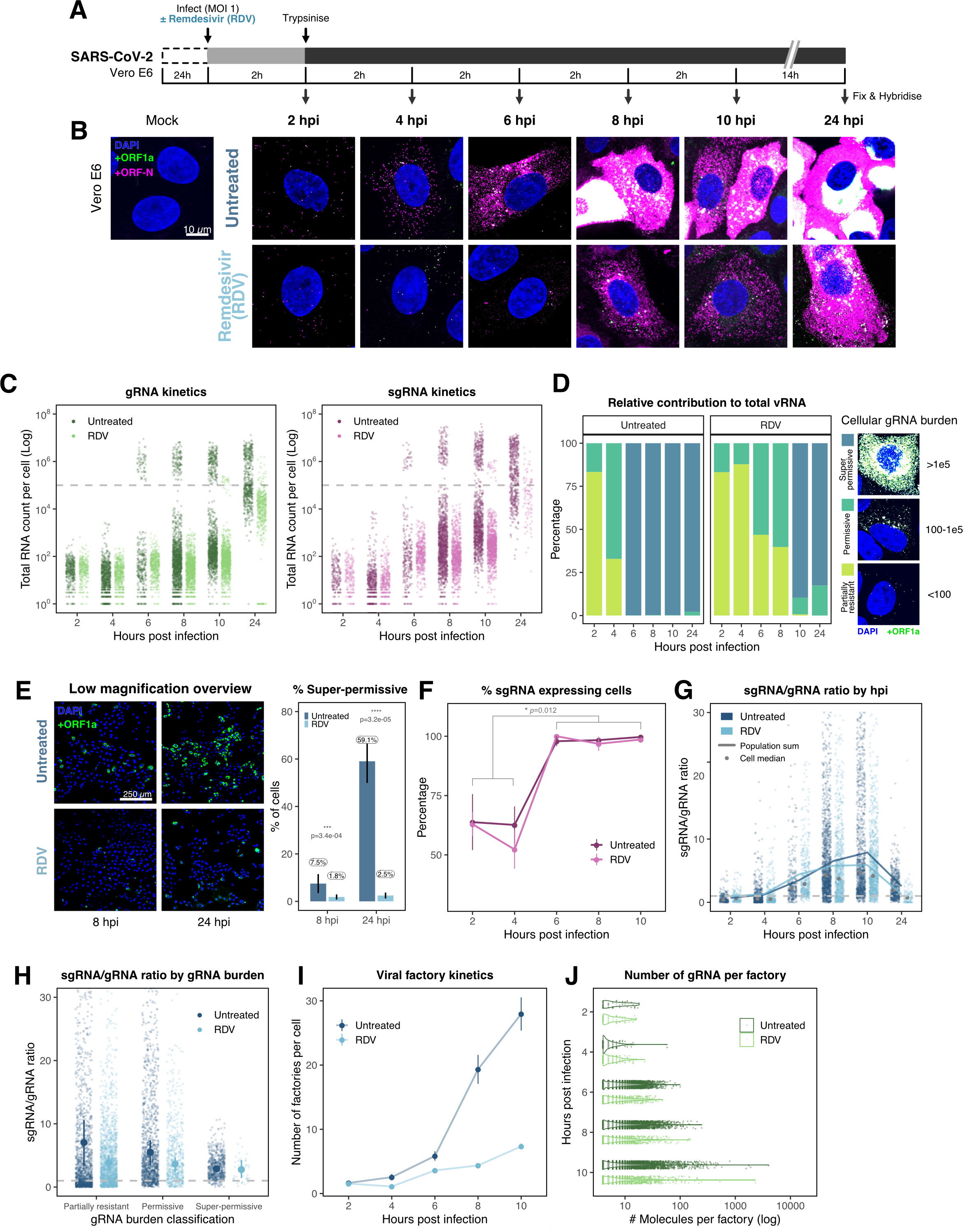
Profiling SARS-CoV-2 replication kinetics at single-molecule resolution. (A) Experimental design to profile SARS-CoV-2 replication kinetics using smFISH. Vero E6 cells were seeded on cover-glass and 24 h later, inoculated with SARS-CoV-2 (VIC strain, MOI = 1) for 2 h. Non-internalised viruses were removed by trypsin digestion and cells fixed at the timepoints shown for hybridisation with +ORF1a and +ORF-N probes. In the remdesivir (RDV) condition, the drug was added to cells at 10 µM during virus inoculation and maintained for the infection period. (B) Maximum z-projected confocal images of infected cells. Numbers at the bottom left corner indicate dynamic contrast range used to display the image. Scale bar = 10 µm. (C) Bigfish quantification of +gRNA or +sgRNA RNA counts per cell. sgRNA counts were calculated by subtracting +ORF1a counts from +ORF-N counts per cell. Horizontal line indicates 105 molecules of RNA. 24 hpi samples and the cells harbouring > 10^7^ RNA counts were quantified by extrapolating single-molecule intensity. Quantified cells from all replicates are plotted. (2-8 hpi, n≥3; 10 and 24 hpi, n=2). Number of cells analysed (Untreated/RDV): 2 hpi, 373/273; 4 hpi, 798/516; 6 hpi, 370/487; 8 hpi, 1442/1022; 10 hpi, 1175/1102; 24 hpi, 542/249. (D) Relative contribution of viral gRNA within the infected cell population. The infected cells were classified into three groups based on gRNA counts: i) ‘partially resistant’ - gRNA<100; ‘permissive’ - 100<gRNA<10^5^; ‘super-permissive’ - gRNA>10^5^. The total gRNA within the infected wells was obtained by summing gRNA counts in population and the figure shows the relative fraction from each classification. Representative max-projected images of cells in each category are shown. (2-8 hpi, n≥3; 10 and 24 hpi, n=2) (E) Identification of super-permissive cells from a low-magnification (20x) high-throughput smFISH assay (left panels). Full z-projected (9 µm) images of infected Vero E6 cells hybridised with +ORF1a probes are shown. At this magnification, only the cells with vRNA count > ∼105 (i.e., corresponding to ‘super-permissive’ cells) exhibit significant smFISH fluorescence compared to uninfected cells. Scale bar = 250 µm. Percentage of super-permissive cells in untreated and RDV-treated conditions at 8 and 24 hpi (right panel). Labels represent average values. Data represented as mean ± S.D. (n=3, ∼ 2000 cells were scanned from each replicate well). Student’s t-test. ***, p<0.001; ****, p<0.0001. (F) Percentage of infected cells expressing sgRNA. sgRNA expressing cells were identified by those having an (+ORF-N – +ORF1a) probe count more than 1. Data represented as mean ± S.E.M. (2-8 hpi, n≥3; 10 and 24 hpi, n=2). (G) Per cell ratio of sgRNA/gRNA counts across the time series. Gray symbols represent cell-to-cell median values whereas the line plot represents ratio calculated from population sum of gRNA and sgRNA. The number of cells analysed are the same as in (Figure 3C), with the exception of cells having equal +ORF1a and +ORF-N probe counts. Horizontal dashed line represents value of 1. (2-8 hpi, n≥3; 10 and 24 hpi, n=2). (H) Per cell ratio of sgRNA/gRNA counts grouped by gRNA burden classification as in (Figure 3D). Data represents median ± S.E.M. Horizontal dashed line represents value of 1. (2-8 hpi, n≥3; 10 and 24 hpi, n=2). (I) The number of viral factories per cell increase over time as assessed by smFISH cluster detection. Cells harbouring > 10^7^ copies of RNA, less than 10 molecules of RNA, cells with no viral factories, and cells from 24 hpi timepoints were excluded from this analysis. Data represented as mean ± S.E.M. Number of cells analysed (Untreated/RDV): 2 hpi, 494/240; 4 hpi, 758/494; 6 hpi, 315/417; 8 hpi, 933/877; 10 hpi, 726/885. (2-8 hpi, n≥3; 10 and 24 hpi, n=2). (J) The kinetics of gRNA copies within viral factories. Spatially extended viral factories were resolved by cluster decomposition to obtain single-molecule counts. The type and number of cells analysed are the same as in (Figure 3I). (2-8 hpi, n≥3; 10 and 24 hpi, n=2).

Our high-throughput FISH intensity analysis showed that this cellular heterogeneity persists beyond the primary rounds of infection. At 24 hpi 40% of the cells in the population did not reach the super-permissive state but formed a distinct population with approximately 10-fold less gRNA (Figure 3C and E). Of note, the high level of gRNA in super-permissive cells (∼10^7^ counts/cell) was comparable throughout the experiment, suggesting an upper limit of vRNA copies in Vero E6 cells (Figure 3C). A similar distribution of gRNA was observed between 24-48 hpi, although the levels of gRNA started to decline, which may reflect cytolytic effects and virus egress (Figure S3B-D). Treating infected cells at 24 hpi with RDV had a modest effect on gRNA levels (Figure S3D) and reduced the cytolytic effects (Figure S3E). We interpret these results as indicating that SARS-CoV-2 +gRNA is long-lived within infected cells and has a long RNA half-life.

Simultaneous analysis of +ORF1a and +ORF-N revealed similar expression kinetics for sgRNA, with 11 copies/cell of sgRNA detected in 63% of infected cells at 2 hpi (Figure 3C and 3F). Since +sgRNA requires -sgRNA template for its production, our results imply that multiple rounds of transcription occur rapidly following virus internalisation that are RDV insensitive. By 6 hpi, most cells contain sgRNA (Figure 3F), with the super-permissive cells supporting high levels of sgRNA transcription. We examined the vRNA replication dynamics and found the ratio of sgRNA/gRNA ranged from 0.5-8 over time (Figure 3G), consistent with a recent report in diagnostic samples (Alexandersen *et al*., 2020). Notably, the sgRNA/gRNA ratio increased between 2 and 10 hpi, followed by a decline at 24 hpi, indicating a shift in preference to produce gRNA over sgRNA in later stages of infection. A similar trend was observed in RDV-treated cells, with a reduced sgRNA/gRNA peak at 8-10 hpi. We estimated the sgRNA/gRNA ratio for individual cells and found that sgRNA synthesis is favoured in the ‘partially resistant’ and ‘permissive’ cells, whereas the ‘super-permissive’ cells had a reduced ratio of sgRNA/gRNA (Figure 3H). In summary, these results indicate that gRNA synthesis is favoured in the late phase of infection, that may reflect the requirement of gRNA to assemble new viral particles.

Positive-sense RNA viruses, including coronaviruses, utilise host membranes to generate viral factories, which are sites of active replication and/or virus assembly (Wolff et al., 2020). Our current knowledge on the genesis and dynamics of these factories in SARS-CoV-2 infection is limited. We exploited the spatial resolution of smFISH to study these structures, which we define as spatially extended foci containing at least 4 gRNA molecules. We observed 1 -2 factories per cell at 2 hpi, which increased to ∼30 factories/cell by 10 hpi (Figure 3I). In addition, the average number of gRNA molecules within these factories, although variable, increased over time (Figure 3J). RDV treatment reduced both the number of viral factories per cell and their RNA content. Together these data show the capability of smFISH to localise and quantify active sites of SARS-CoV-2 replication and to measure changes in gRNA and sgRNA at a single cell level over the course of the infection.

### Super-permissive cells are randomly distributed

Our earlier kinetic analysis of infected Vero E6 cells identified a minor population of ‘super-permissive’ cells containing high gRNA copies at 8 hpi. A random selection of ∼300 cells allowed us to further characterise the infected cell population (Figure 4A-B). To extend these observations we examined the vRNAs in two human lung epithelial cell lines, A549-ACE2 and Calu-3, that are widely used to study SARS-CoV-2 infection (Chu et al., 2020; Hoffmann *et al*., 2020). In agreement with our earlier observations with Vero E6, 3-5% of A549-ACE2 and Calu-3 cells showed a ‘super-permissive’ phenotype (Figure 4C-D). An important question is how these ‘super-permissive’ cells are distributed in the population, as the pattern could highlight potential drivers for susceptibility (Healy et al., 2020). Infection can induce innate signalling that can lead to the expression and secretion of soluble factors such as interferons that induce an anti-viral state in the local cellular environment (Belkowski and Sen, 1987; Schoggins and Rice, 2011). Regulation can be widespread through paracrine signalling or affect only proximal cells. We considered three scenarios where ‘super-permissive’ cells are: randomly distributed, evenly separated or clustered together. We compared the average nearest neighbour distance between ‘super-permissive’ cells and simulated points that were distributed either randomly, evenly or in clusters (Figure S4A-C). In summary, our results show conclusively that the ‘super-permissive’ infected Vero E6, A549-ACE2 and Calu-3 cells were randomly distributed (Figure 4E-F and S4D-E). We interpret these data as being consistent with an intrinsic property of the cell that defines susceptibility to virus infection. The data also argue against cell-to-cell signalling mechanisms that would either lead to clustering (if increasing susceptibility) or to an even distribution (if inhibiting) of infected cells.

**Figure 4.**
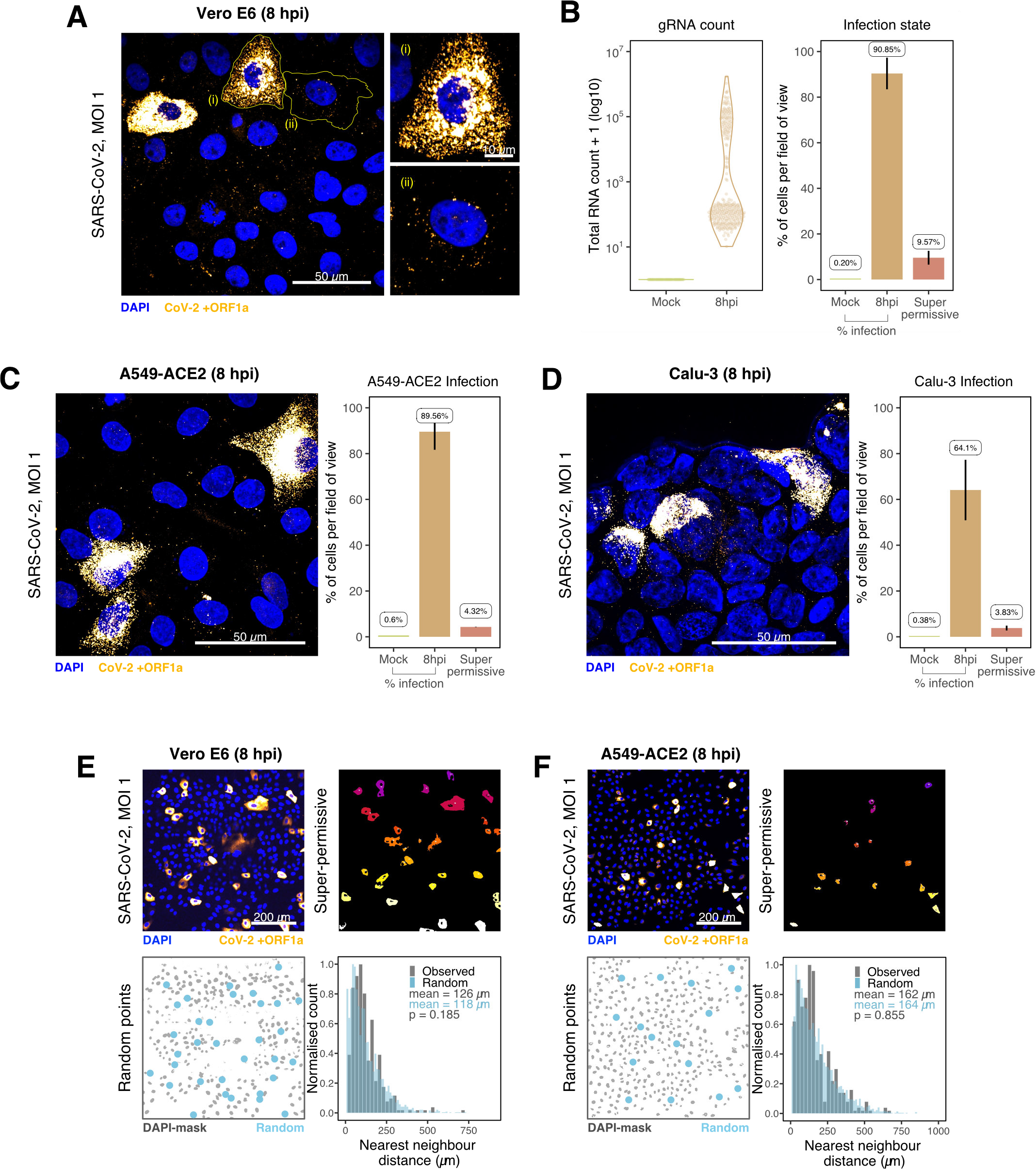
Heterogeneous SARS-CoV-2 RNA replication. (A) Representative 60x magnified field of view (FOV) of SARS-CoV-2 infected Vero E6 cells at 8 hpi (VIC strain, MOI = 1). smFISH against ORF1a was used to visualise cellular heterogeneity in viral RNA counts. Magnified panels show (i) a ‘super-permissive’ cell, and (ii) a cell with discrete viral RNA copies. Scale bar = 10 or 50 µm. (B) Discrete separation of gRNA count distribution among infected cells randomly sampled at 8 hpi, where each dot symbol represents a cell. Statistics for the percentage of infected cells and frequency of ‘super-permissive’ cells at 8 hpi. Quantification was performed per field of view and the number labels represent average values. Cells with > 10^5^ gRNA copies were considered to be ‘super-permissive’, as classified as ‘High’ in (Figure S3C-D) (gRNA quantification: n=4, 148 uninfected and 316 infected cells; Percentage infection: n=3). (C-D) Heterogeneous SARS-CoV-2 replication in lung epithelial A549-ACE2 and Calu-3 cells. The percentage of infected and super-permissive cells were quantified as with Vero E6 cells above. Scale bar = 50 µm (A549-ACE2, n=2; Calu-3, n=3). (E-F) Spatial distribution analysis of super-permissive Vero E6 and A549-ACE2 cells at 8 hpi. Low magnification smFISH overview of infected cells (top left). 2D mask of super-permissive cells (top right). An example of randomly simulated points within the DAPI mask (bottom left). Same number of random points as super-permissive cells were simulated 10 times per FOV. Histogram of nearest neighbour distances calculated from super-permissive cells (Observed) and randomly simulated points (Random) (bottom right). Further modes of spatial analyses are presented in (Figure S4) with infected Calu-3 cells. All confocal images are presented as maximum full z-projection. Data represented as mean ± S.D. Student’s t-test. p-values are shown on the presented visual. (Vero E6, n=3; A549-ACE2, n=2)

### Differential replication kinetics of the B.1.1.7 and Victoria strains

The recent emergence of SARS-CoV-2 VOC, which display differential transmission, pathogenesis and infectivity, have changed the course of the COVID-19 pandemic. Recent studies have focused on mutations in the Spike protein and whether these alter particle uptake into cells and resistance to vaccine or naturally acquired antibodies (Collier et al., 2021; Dicken et al., 2021; Planas et al., 2021). The B.1.1.7 variant is associated with higher transmission (Davies *et al*., 2021; Galloway et al., 2021; Volz *et al*., 2021) and has 17 coding changes mapping to both non-structural (ORF1a/b, ORF3a, ORF8) and structural (Spike and N) proteins. Mutations within the non-structural genes could affect virus replication, independent of Spike mediated entry, thus we used smFISH to compare the replication kinetics of the B.1.1.7 and VIC strains (Figure 5A). We discovered that the number of gRNA molecules at 2 hpi was similar for both viruses, reflecting similar cell uptake of viral particles (Figure 5B-E). However, the quantities of intracellular gRNA and sgRNA were lower in B.1.1.7 infected cells compared to VIC at 6 and 8 hpi (Figure 5E). We also found that while the amount of gRNA per cell was reduced in the B.1.1.7 variant, there were an equal number of +ORF1a and +ORF-N positive cells (Figure 5D), suggesting that the reduced B.1.1.7 RNA burden is due to a differential replication efficiency rather than infection rate. The B.1.1.7 variant also showed a reduced number of replication factories per cell (Figure 5F), with each focus containing on average a lower number of gRNA molecules compared to the VIC strain (Figure 5G). Consistent with the delay in replication, we observed a shallower growth of the sgRNA/gRNA ratio in B.1.1.7 infected cells between 2 to 8 hpi compared to the VIC strain (Figure 5H). These differences between the strains were apparent in all three classifications of cells from our earlier gRNA burden criteria (Figure S5A, 4D). We noted B.1.1.7 infected ‘partially resistant’ and ‘permissive’ cells show lower sgRNA/gRNA ratio while ‘super-permissive’ cells displayed 1.5-fold higher ratio compared to VIC (Figure S5A). The frequency of super-permissive cells was lower for B.1.1.7 at 6 and 8 hpi (Figure 5I and S5B-C) and their distribution was also random (Figure S5D). RDV treatment ablated the differences between the viral strains, demonstrating that the observed phenotype is replication-dependent (Figure 5B, E-I). Nevertheless, the lower level of individual gRNA that we detected in RDV treated cells persisted for at least 8 hpi in both the VIC and B.1.1.7 strains. We conclude that individual gRNA molecules of both the strains are highly stable in the cytoplasm of infected cells.

**Figure 5.**
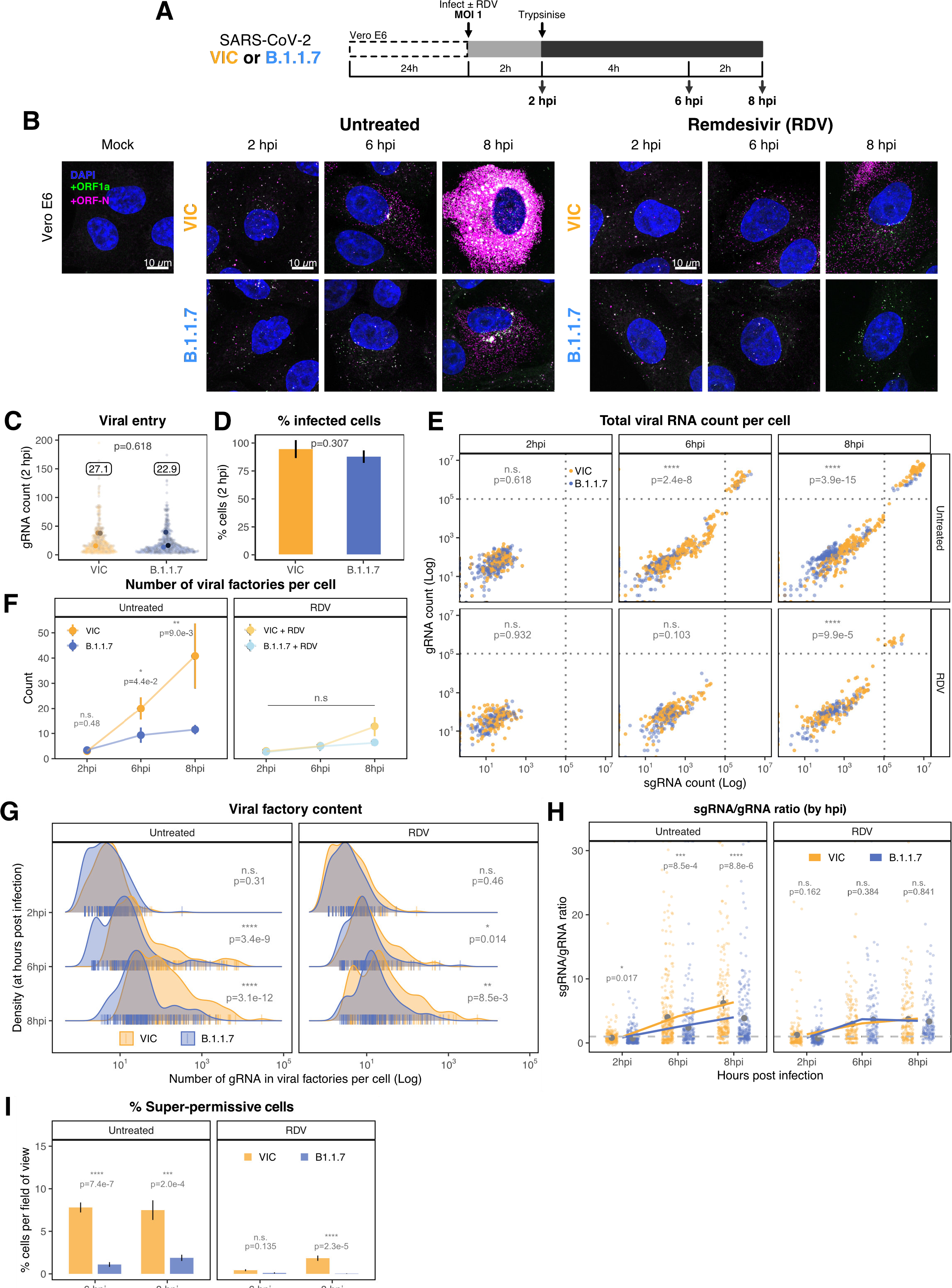
Delayed replication kinetics of B.1.1.7 variant. (A) Experimental design to compare the replication kinetics of VIC and B.1.1.7 SARS-CoV-2 strains. Vero E6 cells were seeded on cover-glass and 24 h later, inoculated with VIC or B.1.1.7 strain (MOI = 1) for 2 h. Non-internalised viruses were removed by trypsin digestion and cells were fixed at designated timepoints for hybridisation with +ORF1a and +ORF-N probes. In remdesivir (RDV) condition, the drug was added to cells at 10 µM during virus inoculation and maintained for the infection period. (B) Maximum z-projected confocal images of Vero E6 cells infected with VIC or B.1.1.7 strains. Representative super-permissive cells from the time series are shown. Scale bar = 10 µm. (C) Comparing viral gRNA counts at 2 hpi between VIC and B.1.1.7. Each symbol represents a cell. Different hue of colours represents readings taken from individual repeat experiments, and the labels represent average values. (n=3; VIC, 424 cells; B.1.1.7, 519 cells). Mann-Whitney U test. (D) Comparing percentage of infected cells between the two viral strains at 2 hpi. Infected cells were determined by +ORF1a smFISH fluorescence. Data represents mean ± S.D. (n=3). Student’s t-test. (E) Bigfish quantification of gRNA and sgRNA smFISH counts per cell. Quantification was performed as in (Figure 3C). Due to bimodality of the data, statistical significance was determined using two-sample Kolmogorov-Smirnov test to compare cumulative distribution of +ORF1a counts between the two strains. (n=3). (VIC, 2 : 6 : 8 hpi = 460 : 343 : 407 cells; B.1.1.7, 2 : 6 : 8 hpi = 396 : 487 : 429 cells). (F) Comparing the number of viral factories per cell between the two viral strains across the time series. Cells harbouring > 10^7^ copies of vRNA were excluded from analysis. Viral factories were identified using Bigfish cluster detection as with (Figure 3I). Data represented as mean ± S.E.M. (n=3). Mann-Whitney U test. (G) Density ridge plot showing the number of gRNA copies within viral factories for VIC and B.1.1.7 variants. The density distribution represents the number of molecules per viral factories per cell. Vertical segment symbol represents a cell. (n=3). Mann-Whitney U test. (H) Per cell ratio of sgRNA/gRNA counts across the time series. Gray symbols represent cell-to-cell mean ± S.E, which are connected by line plots. Horizontal dashed line represents value of 1. (n=3). Mann-Whitney U test. (I) Comparison of the percentage of super-permissive cells between the two strains assessed from low-magnification high-throughput smFISH assay (see (Figure S5A-B) for details). Data represented as mean ± S.D. (n=3). Student’s t-test. n.s., not significant; *, p<0.05; **, p<0.01; ***, p<0.001; ***, p<0.0001.

To test whether our findings using the B.1.1.7 variant are applicable to other cell types, we assessed the replication of both strains in A549-ACE2 cells that were recently reported to be immunocompetent (Li et al., 2021). Both VIC and B.1.1.7 infections resulted in comparable numbers of infected cells and similar numbers of gRNA molecules per cell at 2 hpi, demonstrating a similar degree of viral particle entry into cells (Figure S6A). However, infection with the B.1.1.7 variant led to a reduced gRNA and sgRNA burden at 8 and 24 hpi (Figure 6A-B and S6B-C). Moreover, fewer ‘super-permissive’ cells were detected at these time points (Figure 6C). Hence, the B.1.1.7 variant has also reduced replication rates in an immunocompetent cell line such as A549-ACE2.

**Figure 6.**
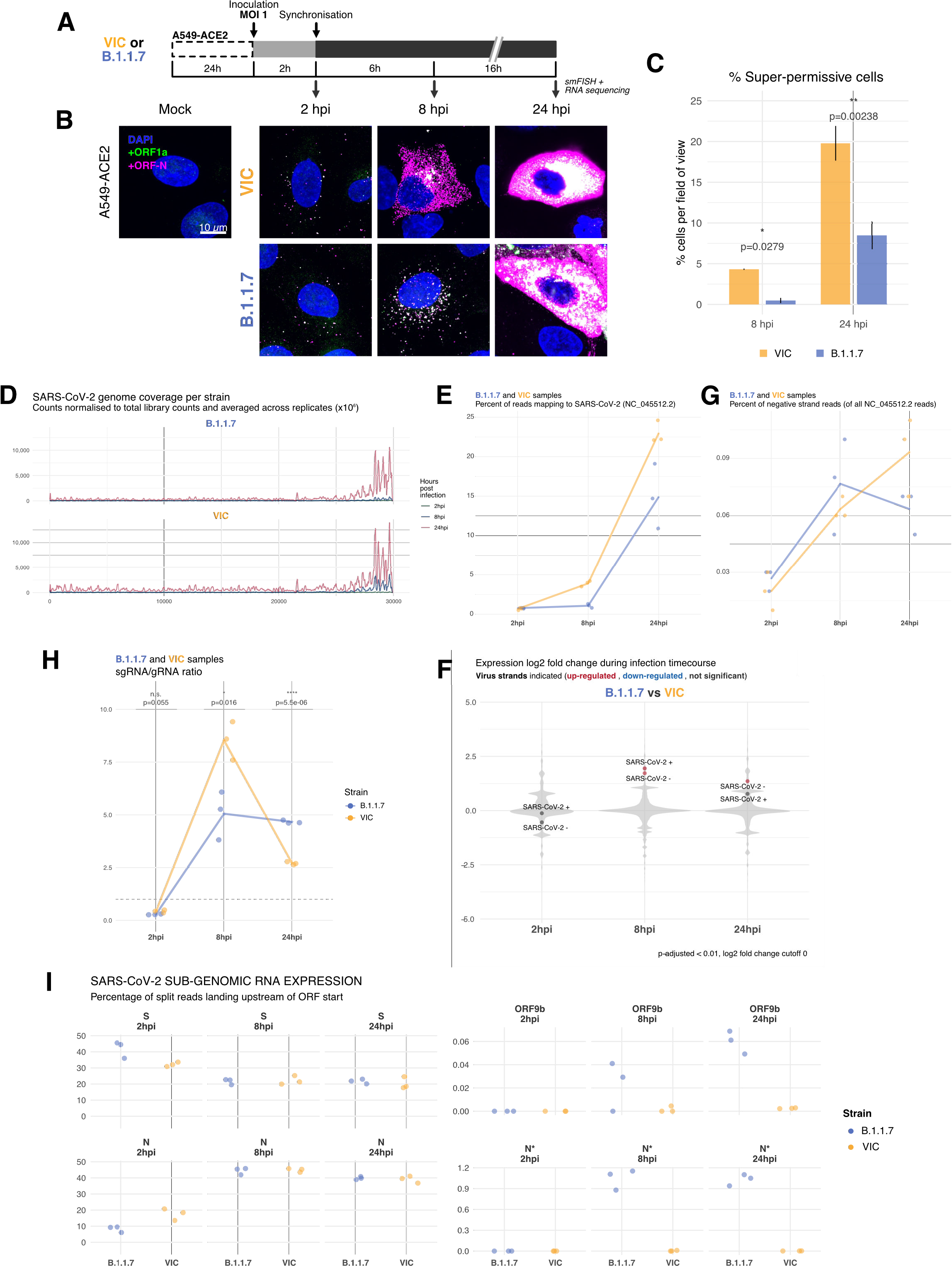
Transcriptomic landscape of B.1.1.7 and Victoria SARS-CoV-2 strains. (A) Experimental design to compare replication kinetics and transcriptomic landscapes of VIC and B.1.1.7 strains. A549-ACE2 cells were seeded and 24 h later, inoculated with VIC or B.1.1.7 strain (MOI = 1) for 2 h. Non-internalised viruses were removed by trypsin digestion and cells were fixed at designated timepoints for smFISH or harvested for RNA-seq library preparation. (B) Maximum z-projected confocal images of A549-ACE2 cells infected with VIC or B.1.1.7. Representative super-permissive cells from the time series are shown. Numbers at the bottom left corner indicate dynamic contrast range used to display the image. Scale bar = 10 µm. (C) Comparison of the percentage of super-permissive cells between the two strains. Super-permissive cells were identified from low-magnification high-throughput smFISH assay (see Figure 7SB-C). Data represented as mean ± S.D. (8 hpi n=2; 24 hpi n=3). *, p<0.5: **, p<0.01. (D) Read coverage along SARS-CoV-2 genome (positive strand) for the two variants in the three timepoints. Counts are normalised to total read count to show the increased proportion of reads from the virus in addition to the accumulation of sub-genomic RNA and averaged across replicates. (n=3). (E) Percentage of reads mapping to SARS-CoV-2 genome of total mapped reads, shown separately for the two strains. Each symbol represents an experimental replicate. (n=3). (F) Violin plots showing fold-changes in the host transcriptome and viral RNA genome comparing B.1.1.7 and VIC strains at the three timepoints. Fold-changes for SARS-CoV-2 positive and negative strands are indicated as separate points and coloured according to the statistical significance of the change (red – higher in VIC, blue – higher in B.1.1.7, gray – no change). p-adjusted < 0.01, log2 fold change cutoff = 0. (n=3). (G) Percent of reads mapping to SARS-CoV-2 negative (antisense) strand relative to all SARS-CoV-2 reads, shown separately for the two strains. Each symbol represents an experimental replicate (n=3). (H) Estimated ratio of SARS-CoV-2 sub-genomic to genomic RNA for the two virus variants at the three timepoints. Student’s t-test. n.s., not significant; *, p<0.05; ****, p<0.001. (n=3). (I) Expression of S, N, ORF9b, and N* viral sub-genomic RNAs in each strain and different timepoints. Expression of each sub-genomic RNA is determined from split reads indicative of transcriptional skipping landing within 100nt upstream of annotated ORF start site, or until upstream ORF start codon if nearer. Percentage of all skip events is shown. (n=3).

To further test our surprising finding regarding reduced replication of the B.1.1.7 with an independent method, we sequenced ribo-depleted total RNA libraries of A549-ACE2 cells infected with VIC or B.1.1.7 for 2, 8 and 24 h (Figure 6A & S6D). As expected, the number of reads mapping to SARS-CoV-2 genome increased over time, reflecting active replication and transcription (Figure 6D). Reads mapping to the 3’ end of the genome increased relative to the 5’ end, reflecting the synthesis of sgRNAs. In agreement with our smFISH analysis, we detected similar levels of vRNA at 2 hpi within B.1.1.7 or VIC infected cells, consistent with similar internalisation rates in A549-ACE2 cells (Figure 6E). However, the abundance of vRNAs in B.1.1.7 infected cells at 8 and 24 hpi was notably lower than with VIC infected cells (Figure 6E). Furthermore, the level of B.1.1.7 RNA was almost unaltered between 2 and 8 hpi, and then increased dramatically at 24 hpi (Figure 6E, S6E), contrasting with VIC infected cells, that showed a continuous increase in vRNA over time. Together, these RNA sequencing data confirm that the B.1.1.7 variant exhibits delayed replication kinetics complementing our smFISH results.

### Transcriptomic changes in B.1.1.7 and Victoria infected cells

To further explore the differences in gene expression between the B.1.1.7 and VIC strains we assessed the abundance of the different vRNAs in infected A549-ACE2 cells. Negative sense viral RNAs represent a small fraction of the vRNA present in the cell, as assayed by smFISH (Figure 6G). These negative sense transcripts are detectable as early as 2 hpi, adding further support to our earlier conclusion that primary viral replication events can occur rapidly post-infection, particularly in ‘super-permissive’ cells (Figure 3C and 6G). The ratio between negative and positive sense vRNAs increased throughout the infection for the VIC strain, but for B.1.1.7 we observed a modest reduction in the ratio at 24 hpi (Figure 6G). To assess the expression of sgRNAs, we quantified the reads mapping to the split junctions derived from RNA-dependent RNA polymerase discontinuous replication (Figure S6F) (Kim *et al*., 2020; V’Kovski *et al*., 2021). In agreement with smFISH data, sgRNAs were detected in low quantities at 2 hpi (Figure 6D and G). For VIC, the sgRNA/gRNA ratio peaks at 8 hpi, followed by a significant drop at 24 hpi (Figure 6G). For B.1.1.7 we observed a significantly lower sgRNA/gRNA ratio at 8 hpi when compared to VIC (Figure 5H). However, the sgRNA/gRNA ratio of B.1.1.7 remained stable between 8 and 24 hpi, surpassing VIC (Figure 6H). These results suggest that both VIC and B.1.1.7 have a different kinetics of gRNA and sgRNA expression, complementing our earlier observations with smFISH (Figure 3C and G)

Next, we assessed the relative abundance of each individual sgRNA. We found that S-sgRNA was the dominant species at 2 hpi, while the sgRNA encoding N (N-sgRNA) become prevalent at 8-24 hpi (Figure S6F). We interpret this early S-to-N sgRNA switch upon infection as indicating a transition to the assembly of viral particles requiring large numbers of N molecules. At 2 hpi B.1.1.7 produced more S-sgRNA but less N-sgRNA than VIC, which is consistent with a delayed B.1.1.7 replication kinetic and S-to-N transition. Furthermore, we found up-regulation of sgRNAs encoding ORF9b (∼0.13%) and N* (∼ 1%) in B.1.1.7 infected cells (Figure 6I), in agreement with recent studies reporting altered sgRNA landscapes for B.1.1.7 (Parker et al., 2021; Thorne et al., 2021). Upregulation of these transcripts is likely to result from advantageous mutations that create novel transcriptional regulatory sequences (TRS-B) in B.1.1.7 (Parker *et al*., 2021; Wang et al., 2021). When we scanned for the TRS motifs in other VOCs and Variants of Interests (VOIs), we found mutations in TRS-B near N* were also found in P.1 (Gamma) and P.2 (Zeta) variants while mutations in TRS-B near ORF9b was unique to B.1.1.7 (see Supplementary Data Item). However, multiple sequence alignment of VOCs and VOIs revealed that mutations accumulate frequently at or near the TRS motif sequences, suggesting SARS-CoV-2 utilises these regulatory motif and surrounding sequences as evolutionary hotspots to modulate sgRNA expression and viral fitness. These transcriptomic results reveal that B.1.1.7 does not only exhibit a delayed replication kinetics, but also produces a differential pool of sgRNAs likely due to mutations within the TRS. Altogether, the novel combination of smFISH and ‘in-bulk’ RNA sequencing that we have described provides a powerful and holistic way to characterise the replication dynamics of SARS-CoV-2. Our pipeline can now be expanded to other VOC, viruses, functional analyses and characterisation of antivirals.

## Discussion

Our spatial quantitation of SARS-CoV-2 replication dynamics at the single molecule and single cell level provides important new insights into the early rate-limiting steps of infection. Typically, analyses of viral replication are carried out using ‘in-bulk’ approaches such as RT-qPCR and conventional RNA-seq. While very informative, these approaches lack spatial information and do not allow single cell analyses. Although single cell RNA-seq analyses can overcome some of these issues (Fiege et al., 2021; Ravindra et al., 2021), their low coverage and lack of information regarding the spatial location of cells remains a significant limitation. In this study we show smFISH is a sensitive approach that allows the absolute quantification of SARS-CoV-2 RNAs at single molecule resolution. Our experiments show the detection of individual gRNA molecules within the first 2h of infection, which most likely reflect incoming viral particles. However, we also observed small numbers of foci comprising several gRNAs sensitive to RDV treatment, demonstrating early replication events. We believe these foci represent ‘replication factories’ as they co-stain with FISH probes specific for negative sense viral RNA and sgRNA. These data provide the first evidence that SARS-CoV-2 replication occurs within the first 2h of infection and increases over time. This contrasts to our observations with the J2 anti-dsRNA antibody where viral dependent signals were apparent at 6 hpi (Cortese *et al*., 2020; Eymieux *et al*., 2021). We noted that co-staining SARS-CoV-2 infected cells with J2 antibody and +ORF1a with an smFISH probe set showed a partial overlap, suggesting that infection may induce changes in cellular dsRNA. These findings highlight the utility of smFISH to uncover new aspects of SARS-CoV-2 replication that are worthy of further study.

We found that SARS-CoV-2 gRNA persisted in the presence of RDV, suggesting a long half-life that may reflect the high secondary structure of the RNA genome that could render it refractory to the action of nucleases (Simmonds et al., 2021). smFISH revealed complex dynamics of gRNA and sgRNA expression that resulted in a rapid expansion of sgRNA (peaking at 8hpi), followed by a shift towards the production of gRNA (24 hpi), results that were confirmed by RNA-seq. Since a viral particle is composed of thousands of proteins and a single RNA molecule, we interpret the high synthesis of sgRNAs as aiming to fulfil the high demand for structural proteins in the viral particles. Once the structural proteins are available in sufficient quantities, the late shift towards gRNA synthesis may ensure the presence of sufficient gRNA to generate the viral progeny.

Our study shows that cells vary in their susceptibility to SARS-CoV-2 infection, where most cells had low vRNA levels (<10^2^ copies/cell), but a minor population (4-10% depending on the cell line) had much higher vRNA burden (>10^5^ copies/cell) at 10 hpi. In contrast, the number of intracellular vRNA copies at 2 hpi was similar across the culture, suggesting that this phenotype is not explained by differences in virus entry. These ‘super-permissive’ cells account for the majority of vRNA within the culture and mask the dominant cell population. Similar results were obtained with Vero E6, Calu-3 and A549-ACE2 suggesting this is a common feature of SARS-CoV-2 infection. As both Calu-3 and A549-ACE2 have intact innate sensing pathways (Cao et al., 2021; Li *et al*., 2021) unlike Vero cells (Desmyter et al., 1968), this variable susceptibility is unlikely to reflect differential immune cell signalling and is consistent with their random distribution within the culture. The reason for the differential infection fitness may rely on the intrinsic properties of each cell, including the stage of the cell cycle, the expression of individual antiviral sensors or the metabolic state. Recent single cell RNA sequencing studies of SARS-CoV-2 infected bronchial cultures identified ciliated cells as the primary target. However, only a minority of these cells contained vRNA that may either reflect low sequencing depth or cell-to-cell variation in susceptibility (Fiege *et al*., 2021; Ravindra *et al*., 2021). The human respiratory tract encompasses the nasal passage, large and small airways and bronchioles and our knowledge on how specific cell types and SARS-CoV-2 RNA burden relate is still limited. Applying smFISH to clinical biopsies and experimentally infected animal samples (Salguero et al., 2021) will allow us to address this important question.

Given the current status of the pandemic, there has been a global effort to understand the biology of emergent VOC with high transmission rates and possible resistance to neutralising antibodies. Most studies have focused on mutations mapping to the Spike glycoprotein as they can alter virus attachment, entry and sensitivity to vaccine induced or naturally acquired neutralising antibodies. However, many of the mutations map to other viral proteins, including components of the RNA-dependent RNA polymerase complex that could impact RNA replication, and non-coding regulatory regions as the TRSs, which can affect sgRNA expression. Our smFISH analysis revealed that the B.1.1.7 variant shows slower replication kinetics compared to the VIC strain, resulting in lower gRNA and sgRNA copies per cell, fewer viral replication factories and a reduced frequency of ‘super-permissive’ cells. This delay in B.1.1.7 replication was observed in Vero and A549-ACE2 cells and was confirmed by RNA-seq as an orthogonal method.

Emerging VOC, such as B.1.1.7, have been reported to have a fitness advantage in terms of their ability to transmit compared to the VIC isolate (Caly *et al*., 2020; Davies *et al*., 2021; Kidd *et al*., 2021; Volz *et al*., 2021). However, the mechanisms underlying increased transmission are not well understood. Interestingly, a recent study reported that B.1.1.7 leads to higher levels of intracellular vRNA and N protein than VIC at 24 and 48 hpi using ‘in-bulk’ RT-qPCR and immunofluorescence, respectively (Thorne *et al*., 2021). We observed that while B.1.1.7 still produces lower level of vRNA than VIC at 24 hpi (Figure 6E), it exhibits a clear recovery compared to 8 hpi. It is thus plausible that both variants yield similar total amounts of viral RNA and proteins but within a different time frame. The potential differences in replication dynamics between the two variants are also reflected in distinct sgRNA/gRNA ratios throughout the infection (Figure 6H). That said, ‘in-bulk’ RT-qPCR analysis does not provide absolute quantification and individual cell assessment and, therefore, should likely be biased towards the super-susceptible cells that account for most of the RNA burden. Thorne and colleagues also reported an elevated expression of the sgRNA encoding the innate agonist ORF9b (Thorne *et al*., 2021), which is also supported by our results. We noticed that the increase of ORF9b sgRNA expression may be due to mutations in non-coding regulatory sequences involved in discontinuous replication (TRS), and that such mutations are common across VOCs possibly mediating differential sgRNA expression. Enhanced ORF9b expression, together with the lower intracellular vRNA levels present in B.1.1.7 infected cells, may grant this variant with an advantage to evade the antiviral response. This advantage combined with mutations in the Spike that are proposed to improve cell entry could provide the B.1.1.7 with a replicative advantage over the early lineage VIC strain enabling its rapid dissemination across the human population (Caly *et al*., 2020; Davies *et al*., 2021; Kidd *et al*., 2021; Volz *et al*., 2021). A recent longitudinal study of nasopharyngeal swabs showed that the B.1.1.7 variant was associated with longer infection times and yet showed similar peak viral loads to non-B.1.1.7 variants (Kissler *et al*., 2021). The authors conclude that this extended duration of virus shedding may contribute to increased transmissibility and is consistent with our data showing reduced replication of B.1.1.7 at the single cell level. Replication fitness will be defined by the relationship of the virus with its host cell, aggressive replication is expected to trigger cellular antiviral sensors. In contrast, lower replication may allow the virus to replicate and persist for longer periods before host antiviral sensors are triggered. Such differences, and their impact on host anti-viral responses, are likely to be of key importance for our understanding of the success of viral variants to spread through the population.

## STAR★Methods

### Cell culture

Vero E6, A549-ACE2 (kind gift from the Bartenschlager lab) (Klein *et al*., 2020) and Huh-7.5 cells were maintained in standard DMEM, Calu-3 cells in Advanced DMEM both supplemented with 10% fetal bovine serum, 2mM L-glutamine, 100 U/mL penicillin and 10μg/mL streptomycin and non-essential amino acids. All cell lines were maintained at 37°C and 5% CO2 in a standard culture incubator.

### Virus propagation and infection

SARS-CoV-2 strains: Victoria 01/20 (BVIC01) (Caly *et al*., 2020) (Provided by PHE Porton Down after supply from the Doherty Centre Melbourne, Australia) and B.1.1.7 (Tegally et al., 2020) (20I/501Y.V1.HMPP1) (provided by PHE Porton Down). Viral strains were propagated in Vero E6 cells as described (Wing et al., 2021). Briefly, naïve Vero E6 cells were infected with SARS-CoV-2 at a multiplicity of infection (MOI) of 0.003 and incubated for 48-72h until visible cytopathic effect was observed. At this point, cultures were harvested, clarified by centrifugation to remove residual cell debris and stored at -80oC. To determine the viral titre, fresh Vero E6 cells were inoculated with serial dilutions of SARS-CoV-2 viral stocks for 2h followed by addition of a semi-solid overlay consisting of 1.5% carboxymethyl cellulose (SIGMA). Cells were incubated for 72h and visible plaques enumerated by fixing cells using amido black stain to calculate plaque-forming units (PFU) per ml. Similarly, HCoV-229E (Andrew Davidson lab (Bristol) and Peter Simmmonds lab (Oxford)) virus was propagated in Vero E6 cells and TCID50 was performed in Huh-7.5 cells.

For smFISH experiments with the SARS-CoV-2 stains, cells were infected at an MOI of 1 for 2h followed by extensive washing in PBS. Residual cell surface associated virus was removed by trypsin treatment of the cell monolayer for 2mins followed by neutralisation of the trypsin using serum containing media. Infected cells were then maintained for defined periods up to 24h. For the HCoV-229E, cells were infected at an MOI of 1 and were maintained for 24 and 48h.

### RT-qPCR

Infected cells were harvested in RLT buffer and RNA extracted using the Qiagen RNAeasy kit. SARS-CoV-2 RNA was quantified using a one-step reverse transcriptase qPCR (RT-qPCR) kit (Takyon) in a multiplexed reaction containing primer probes directed against the SARS-CoV-2 N gene (FAM) and ß-2-Microglobulin (VIC) as an internal control. All qPCR reactions were carried out using a Roche 96 Light cycler (Roche). (SARS primer probe IDT CAT:100006770, B2M primer probe Applied Biosystems 4325797)

### Single molecule fluorescence in situ hybridisation (smFISH)

smFISH was carried out as previously reported (Titlow *et al*., 2018; Yang et al., 2017) with minor modifications. Briefly, cells were grown on #1.5 round glass coverslips in 24-well plate or in µ-Slides 8 well glass bottom (IBIDI) and fixed in 4% paraformaldehyde (Thermo Fisher) for 30 min at room temperature. Cells were permeabilised in PBS/0.1% Triton X-100 for 10 min at room temperature followed by washes in PBS and 2x SSC. Cells were pre-hybridised in pre-warmed (37°C) wash solution (2x SSC, 10% formamide) twice for 20 min each at 37°C. Hybridisation was carried out in hybridisation solution (2x SSC, 10% formamide, 10% dextran sulphate) containing 500 nM smFISH probes overnight at 37°C. For infection timepoints beyond 24 h, smFISH probes were added at 1 µM. After the overnight hybridisation, cells were washed for 20 min in pre-warmed wash solution at 37°C followed by counterstain with DAPI (1 µg/ml), phalloidin-Alexa Fluor 488 conjugate (264 nM) and/or CellMask Green (1:1,000,000) diluted in wash solution. Cells were then washed once with wash solution for 20 min at 37°C and twice with 2x SSC for 10 min each at room temperature. Cells were mounted using Vectashield, IBIDI mounting media or 2x SSC.

For RNase digestion experiments, RNaseT1 (Thermo Fisher, EN0541, 100 U/ml) or RNaseIII (M0245S, NEB, 20 U/ml) was used to degrade single-stranded RNA and double-stranded RNA, respectively. Permeabilised cells were treated with RNases in PBS supplemented with 5 mM MgCl_2_ and incubated at 37°C for 1 h and washed three times with PBS.

In the experiment to detect viral negative strands, double-stranded RNA (dsRNA) was denatured using DMSO, formamide or NaOH. After the permeabilisation step, cells were rinsed in distilled water and were treated with 50 mM NaOH for 30s at room temperature, 70% formamide at 70°C for 1 h or 90% DMSO at 70°C for 1 h. Following the treatments, cells were quickly cooled on ice, washed in ice-cold PBS and subjected to standard smFISH protocol. The smFISH experiments in Figure 3 and Figure 5 were performed with DMSO and heat denaturation.

### smFISH probe design and specificity analysis

Candidate smFISH probe sequences were acquired using Stellaris® Probe Designer version 4.2 (https://www.biosearchtech.com/stellaris-designer) with following parameters: Organism – Human, Masking level – 5, Oligo length – 20 nt, Minimum spacing length – 3 nt. Appropriate region of the SARS-CoV-2 Wuhan-Hu-1 (NC_045512.2) reference sequence was used as target sequence. Candidate sequences were BLAST screened against custom human transcriptome and intron database to score number of off-target base-pair matches, then 35 – 48 sequences with the least match scores were chosen per probe set. Oligonucleotides were singly labelled with ATTO633, ATTO565, Cy3, or ATTO488 at 3’ ends according to a published protocol (Gaspar *et al*., 2017) and were concentration normalised to 25 µM. All probe sets used in this study had degree of labelling > 0.94.

For in silico probe sequence specificity analysis, selected oligonucleotide sequences were aligned against SARS-CoV-1 (NC_004718), SARS-CoV-2 (NC_045512), MERS-CoV (NC_019843), HCoV-229E (NC_002645), HCoV-NL63 (NC_005831), HCoV-OC43 (NC_006213), HCoV-HKU1 (NC_006577), Human (GCF_000001405.39), and African green monkey (GCF_015252025.1) RefSeq genome or transcriptome assembly using ‘bowtie2’ (2.4.4) (Langmead and Salzberg, 2012). Following bowtie2 arguments were used to find minimum edit distance of oligonucleotide sequences to target genome/transcriptome: --end-to-end --no-unal --align-seed-mm 0, --align-seed-length 5, --align-seed-interval 1-1.15, -- effort-extend 15, --effort-repeat 2. Melting temperatures were obtained using ‘rmelting’ (1.8.0) R package at 300mM Na concentration (2x SSC). smFISH probe sequences used in this study are available in Supplementary Table S1.

### Immunofluorescence

After permeabilisation, cells were blocked in blocking solution (50% Li-Cor Odyssey blocking solution, pretreated with RNASecure for 30 min and supplemented with 2 mM ribonucleoside vanadyl complex and 0.1% Tween-20) for 30 min at room temperature. Then, cells were incubated with J2 primary antibody (Scicons 10010200) at 0.5 µg/ml or human anti-N primary antibody (Ey2B clone 1:2000) (Huang et al., 2020) for 2 h at room temperature. Cells were washed three times in PBS/ 0.1% Tween-20 (PBSTw) for 10 min each at room temperature and incubated with fluorescent secondary antibodies (1:500) diluted in blocking solution for 1 h at room temperature. After further three washes in PBSTw, cells were mounted using Vectashield or IBIDI mounting media. For combined smFISH and immunofluorescence, antibody staining was carried out sequentially after the smFISH protocol.

### Microscopy and image handling

Cells were imaged on an Olympus SpinSR10 spinning disk confocal system equipped with Prime BSI and Prime 95B sCMOS cameras. Objectives used were 20x dry (0.8 NA, UPLXAPO20X), 60x silicone oil (1.3 NA, UPLSAPO60XS2), 60x oil (1.5 NA, UPLAPOHR60X), or 100x oil (1.45 NA, UPLXAPO100XO). Image voxel sizes were 0.55 x 0.55 x 2 µm (x:y:z) with the 20x objective and 0.11 x 0.11 x 0.2 µm (x:y:z) with the 60x and 100x objectives. Automatic and manual image acquisition and image stitching were performed with Olympus cellSens Dimension software. Images were uploaded and stored in the University of Oxford OMERO server (Allan et al., 2012) and OMERO.figure (3.2.0) was used to generate presented image visualisations.

## Image analysis

### Cell segmentation and counting

Cell segmentation was performed either manually in ImageJ (National Institute of Health) or automatically with Cellpose (0.6.1) (Stringer et al., 2021) using 2D maximum intensity projected images of phalloidin or CellMask stains. Cellpose parameters for 60x and 100x magnification images were model_type=cyto, diameter=375, flow_threshold=0.9, cellprob_threshold=-3. For 20x stitched images, CellMask channel was deconvolved with constrained iterative module using cellSens (5 iterations, default spinning disk PSF, Olympus), then following Cellpose parameters were used: model_type=cyto, diameter=55, flow_threshold=0, cellprob_threshold=-6. Total number of cells per image was counted using a custom ImageJ macro script or from the Cellpose segmentation output on DAPI channel images (model_type=nuclei, diameter=20, default threshold). Infected cells were counted using ImageJ “3D object counter” or manually.

### Quantification of smFISH images

Single-molecule level quantification of smFISH images were performed either with FISH-quant (Mueller et al., 2013) or Bigfish (https://github.com/fish-quant/big-fish). For FISH-quant, ImageJ ROI files were converted to FQ outline file using a custom python script. Then, smFISH channels were Laplacian of Gaussian filtered (sigma = 7, 3 px) and pre-detected using local maximum mode with “allow smaller z region for analysis” option enabled. Pre-detected diffraction limited spots were fitted with 3D Gaussian and thresholded in batch mode based on filtered intensity, amplitude and σz. Thresholds were defined by uninfected “Mock” condition samples. Viral factories were quantified using the TxSite quantification mode of FISH-quant (xy:z = 500:1200 nm crop per factory) with normal-sampled averaged single-molecule image (xy:z = 15:12 px) from batch mode output by integrated intensity in 3D method. Similarly, cells displaying high viral load were quantified by analogously integrating smFISH intensities, subtracting uninfected condition background fluorescence and dividing by the intensity of reference single-molecules.

Large smFISH datasets were processed with a custom python pipeline using Bigfish, skimage, and numpy libraries (available in the Github repository). Tif files were converted to a numpy array and individual cells were segmented from the image using the Cellpose library as described above. Images where cells were labelled with the CellMask stain were pre-processed with a median filter, radius = 50. Background signal in the smFISH channel was subtracted with the skimage.white_tophat algorithm (radius = 5, individual z frames were processed in 2D due to memory constraints, results were indistinguishable from 3D-processed images). Threshold setting for smFISH spot detection was set specifically for each set of images collected in each session. Viral factories were resolved using ‘decompose_cluster()’ function to find a reference single-molecule spot in a less signal-dense region of the image, which was used to simulate fitting of the gaussian modelised reference spot into viral factories until the local signal intensities were matched. Decomposed spots were grouped into clusters with previously reported radii of double-membrane vesicles (DMV) measured by electron microscopy (150nm pre-8hpi and 200nm post-8hpi) (Cortese *et al*., 2020).

### Dual colour smFISH spot detection analysis

The same viral RNA target was detected using two smFISH probes labelled with alternating (ODD and EVEN) red and far-red fluorochromes. Resulting images were processed in FISH-quant to obtain 3D coordinates of each spots. Percentage co-localisation analysis was performed with a custom script using an R package “FNN” (1.1.3). Briefly, we calculated 3D distance of nearest neighbour for each spot in the red channel to the closest detected spot in the other channel and repeated the analysis starting from the far-red channel. We then used a value of 300 nm to define co-localised spots corresponding to the same viral RNA molecule. The presented visuals report percentage co-localisations calculated from the red channel to the far-red channel and vice-versa. The analysis was performed per field of view.

### Quantification of fluorescence intensity and signal colocalization

Immunofluorescence images were background subtracted using rolling ball subtraction method (radius = 150 px) in ImageJ. Anti-dsRNA (J2) stain was quantified by integrating fluorescence signal across the z-stacks of cellular region of interest divided by the cell volume to obtain signal density. Signal density was normalised to the average signal density of uninfected “Mock” condition cells. Fluorescence intensity profiles were obtained using ImageJ “plot profile” tool across 3 µm region on 1 µm maximum intensity projected images. To assess colocalisation of N protein with SARS-CoV-2 RNA, ellipsoid mask centred around centroid xyz coordinates of smFISH spots were generated with the size of the point-spread function (xy radius=65 nm, z radius=150 nm) using ImageJ 3D suite. Integrated density of N-protein channel (background subtracted, radius=5px) fluorescence within the ellipsoid mask was measured and compared to the equivalent signal in the uninfected condition.

### Simulation of highly permissive cell distribution

Simulations were performed to determine if the appearance of SARS-CoV-2 super-permissive cells follows a random distribution. The general strategy was to test the complete spatial randomness hypothesis by comparing the average nearest neighbour distance of superinfected cells to an equal number of randomly selected coordinates (Ripley, 1979). 2D spatial coordinates of superinfected cells were obtained from the 3D-object counter (ImageJ) as described above. Cell nuclei were segmented with the DAPI channel and placement of random coordinates was confined to pixels that fell within the DAPI segmentation mask. Nearest neighbour distances were calculated using the KDtree algorithm (Maneewongvatana and Mount, 1999) implemented in python (scipy.spatial.KDTree). Pseudo-random distributions were simulated by randomly placing the first coordinate, then constraining the placement of subsequent coordinates to within a defined number of pixels.

### RNA-sequencing library preparation

RNA from infected cells were extracted as described above. Sequencing libraries were prepared using the Illumina Total RNA Prep with Ribo-Zero Plus library kit (Cat# 20040525) according to manufacturer’s guidelines. Briefly, 100ng of total RNA was first depleted of the abundant ribosomal RNA present in the samples by rRNA targeted DNA probe capture followed by enzymatic digestion. Samples were then purified by Beckman Coulter RNAClean XP beads (Cat #A63987). Obtained rRNA-depleted RNA was fragmented, reverse transcribed, converted to dsDNA, end repaired and A-tailed. The A-tailed DNA fragments were ligated to anchors allowing for PCR amplification with Illumina unique dual indexing primers (Cat#20040553). Libraries were pooled in equimolar concentrations and sequenced on Illumina NextSeq 500 and NextSeq 550 sequencers using high-output cartridges (Cat# 20024907), generating single 150nt long reads.

## RNA-sequencing analysis

### Genomes

We downloaded the human genome primary assembly and annotation from ENSEMBL (GRCh38.99) and the SARS-CoV-2 RefSeq reference genome from NCBI (NC_045512.2). We combined the human and viral genome and annotation files into one composite genome and annotation file for downstream analyses.

### Alignment and gene counts

We performed a splice-site aware mapping of the sequencing reads to the combined human and SARS-CoV-2 genome and annotation using STAR aligner (2.7.3a) (Dobin et al., 2013). We also used STAR to assign uniquely mapping reads in strand-specific fashion to the ENSEMBL human gene annotation and the two SARS-CoV-2 strains.

### Principal Component Analysis

To assess if SARS2 infection is the main driver of differences in the RNA-seq samples, we performed a principal component analysis (PCA). First, we performed library size correction and variance stabilisation with regularized–logarithm transformation implemented in DESeq2 (1.28.1) (Love et al., 2014). This corrects for the fact that in RNA-seq data, variance grows with the mean and therefore, without suitable correction, only the most highly expressed genes drive the clustering. We then used the 500 genes showing the highest variance to perform PCA using the prcomp function implemented in the base R package stats (4.0.2) (R Core Team 2020).

### Differential Expression Analysis

We performed differential expression analysis using the R package DESeq2 (1.28.1) (Love *et al*., 2014). DESeq2 estimates variance-mean dependence in count data from high-throughput sequencing data and tests for differential expression based on a model using the negative binomial distribution.

### SARS-CoV-2 sub-genomic RNA expression

To assess relative levels of viral sub-genomic and genomic RNA expression, we tallied the alignments (using GenomicRanges and GenomicAlignments R packages (Lawrence et al., 2013)) mapping to the region unique to the genomic RNA and the shared region and normalised for their respective lengths. Unique contribution of sgRNA region was then estimated by subtracting the contribution of the genomic RNA from the shared region. In order to assess expression of individual SARS-CoV-2 sub-genomic RNAs, we extracted split (junction) reads mapping to the viral genome with the GenomicAlignments R package (1.24.0) (Lawrence *et al*., 2013). The sub-genomic transcripts fully overlap the full genomic RNA, and partially with each other. While the molecular process generating these sub-genomic RNAs is distinct from RNA splicing, from the point of view of short read mapping they are equivalent. We determined the relative expression level of each sgRNA generated by transcriptional skipping by calculating the number of reads supporting skipping into a region upstream of each annotated viral ORF. To avoid spurious mappings, we filtered for skip sites that were present in all three replicates and constituted at least 0.1% of all skipped viral reads.

### Statistics, data wrangling and visualisation

Statistical analyses were performed in R (3.6.3) and RStudio (1.4) environment using an R package “rstatix” (0.7.0). P values were adjusted using the Bonferroni method for multiple comparisons. The “tidyverse suite” (1.3.0) was used in R, and “Numpy” and “Pandas” python packages were used in Jupyter notebook for data wrangling. Following R packages were used to create the presented visualisation: “ggplot2” (3.3.2), “ggbeeswarm” (0.6.0), “hrbrthemes” (0.8.0), “scales” (1.1.1), and “patchwork” (1.1.1). Further visual annotations were made in the Affinity Designer (Serif).

## Supplemental information

### Supplementary figures

Supplementary Figure S1-6 and legends.

Supplementary data item

Multiple sequence alignment of SARS-CoV-2 variants of concern and variants of interest centred around transcriptional regulatory sequence motifs.

Supplementary tables

Table S1. smFISH oligonucleotide probe sequences used in this study.

Table S2. Differentially expressed gene analysis of RNA sequencing samples from A549-ACE2 cells infected with B.1.1.7 or Victoria SARS-CoV-2 strains at 2, 8 and 24 hours post infection.

## Author contributions

J.Y.L. designed and conducted experiments, designed and carried out quantitative image and bioinformatic analysis pipelines, drafted the figures and co-wrote the manuscript. P.A.C.W designed and conducted experiments and co-revised the manuscript. D.S.G. conducted experiments and quantitative image analysis. M.N. carried out experiments and set up virus infection in Category III containment labs. A.I.J. designed and carried out bioinformatics and next-generation sequencing analysis. J.T. designed and conducted quantitative image analysis of viral RNA numbers and distribution. X.Z. carried out experiments (in Category III containment labs). N.P. carried out sequencing experiments (on infection time courses of variants). L.I. provided resources (Coronavirus strain causing human cold). M.K.T. designed and wrote quantitative software for viral RNA stability. R.M.P. provided advice and discussion for advanced microscopy performance and image analysis. A.W. provided resources (set up bespoke microscopes and labs for this project, and quality control of microscope performance). D.D.A. provided discussion of the design of the experiments and their interpretation. W.J. provided resources (the Alpha variant and facilitated the extensive and complex Containment Category III work at the Sir William Dunn School of Pathology). A.C. co-designed the study, the bioinformatics analysis methods and co-wrote the manuscript. J.A.M. co-designed the study and co-wrote the manuscript. I.D. co-designed the study, the computational methods for quantitative analysis of virus numbers, single cell analysis and spatial distributions, and co-wrote the manuscript.

## Supporting information

Supplemental Data Item

## Acknowledgements

We are grateful to Danail Stoychev and Maria Kiourlappou for the advice on Python programming and high-performance computing. We are very grateful to Olympus UK and Europe for their generous loan of an Olympus IXplore SpinSR spinning disk system for the imaging work in this project and to Matthew Freeman and Jordan Raff for enabling us to install the microscope in the Dunn School of Pathology specifically for this SARS-CoV-2 work. We thank Michael Knight, Maeva Dupont, Lisa Chauveau, and Javier Gilbert Jaramillo for their provision of resources and assistance in Category III containment labs. We thank Micron Advanced Imaging Unit (https://micronoxford.com) for provision of advanced microscopy facilities and technical advice.

## Supplementary figure legends

**Figure S1.**
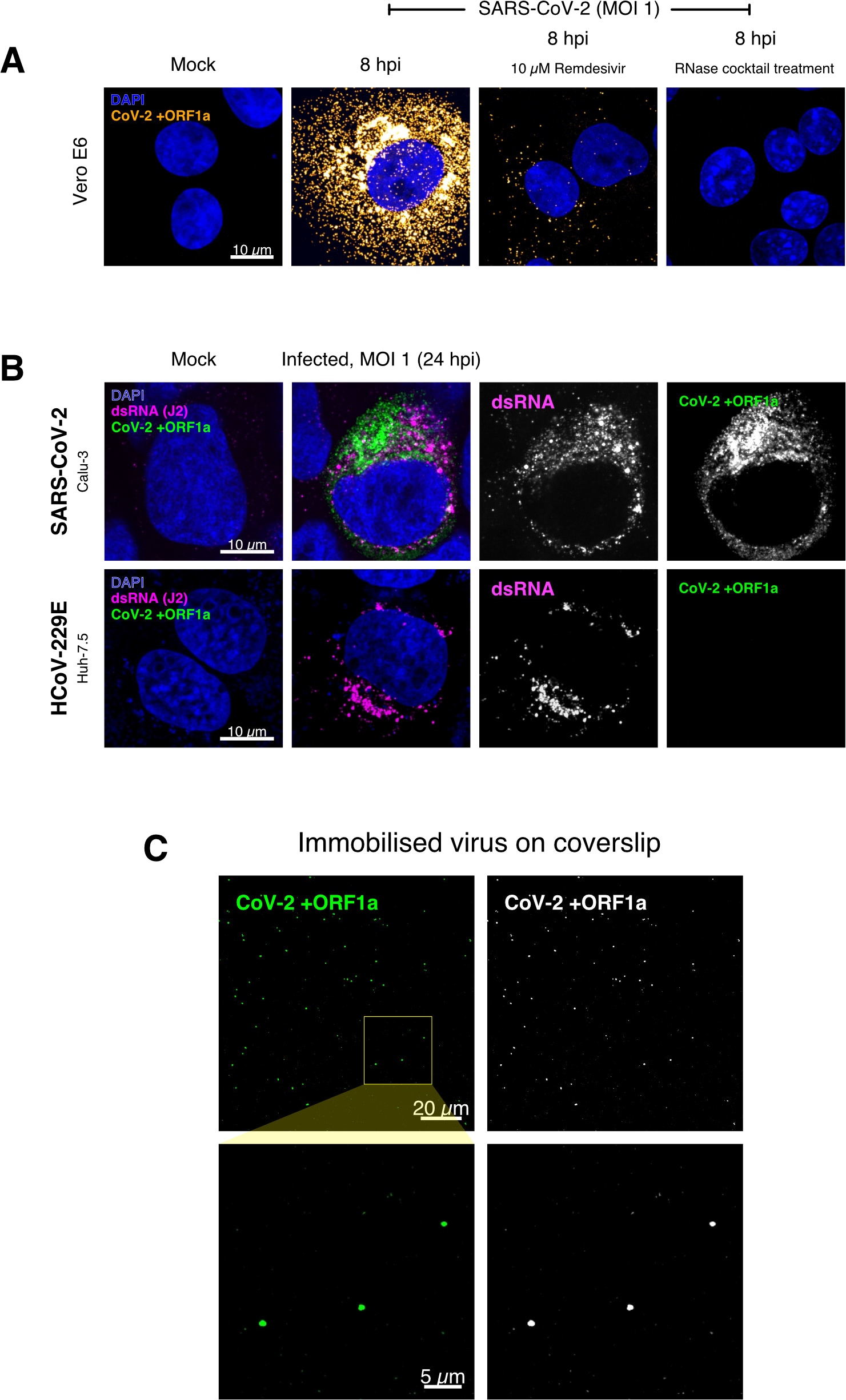
Specific detection of SARS-CoV-2 RNA using single-molecule fluorescence in situ hybridisation (smFISH), related to Figure 1. (A) Specificity of the +ORF1a smFISH probe for SARS-CoV-2 RNA. Vero E6 cells were infected with SARS-CoV-2 (VIC, MOI = 1), fixed at 8 hpi and hybridised with +ORF1a smFISH probe. In the remdesivir (RDV) condition, the drug was added to the cells at 10 µM during virus inoculation and maintained for the infection period. For the RNase digestion, permeabilised cells were treated with a cocktail of RNaseT1 and RNaseIII in the presence of MgCl_2_ to digest RNA prior to probe hybridisation. Representative full z-projection (8 µm) confocal images are shown. Scale bar = 10 µm. (B) Calu-3 (top panels) and Huh-7.5 (lower panels) cells were infected with SARS-CoV-2 (VIC) and HCoV-229E (MOI = 1), respectively, fixed at 24 hpi and hybridised with the SARS-CoV-2-specific +ORF1a probe. In addition, cells were stained with anti-dsRNA (J2) to identify heavily infected cells. Representative single slice confocal images are shown. Scale bar = 10 µm. (C) Visualisation of encapsidated SARS-CoV-2 RNA with smFISH. Virus was immobilised onto poly-L-lysine-coated coverslips and visualised via the +ORF1a probe. A 1 µm maximum z-projected confocal image is shown. Scale bar = 20 µm or 5 µm.

**Figure S2.**
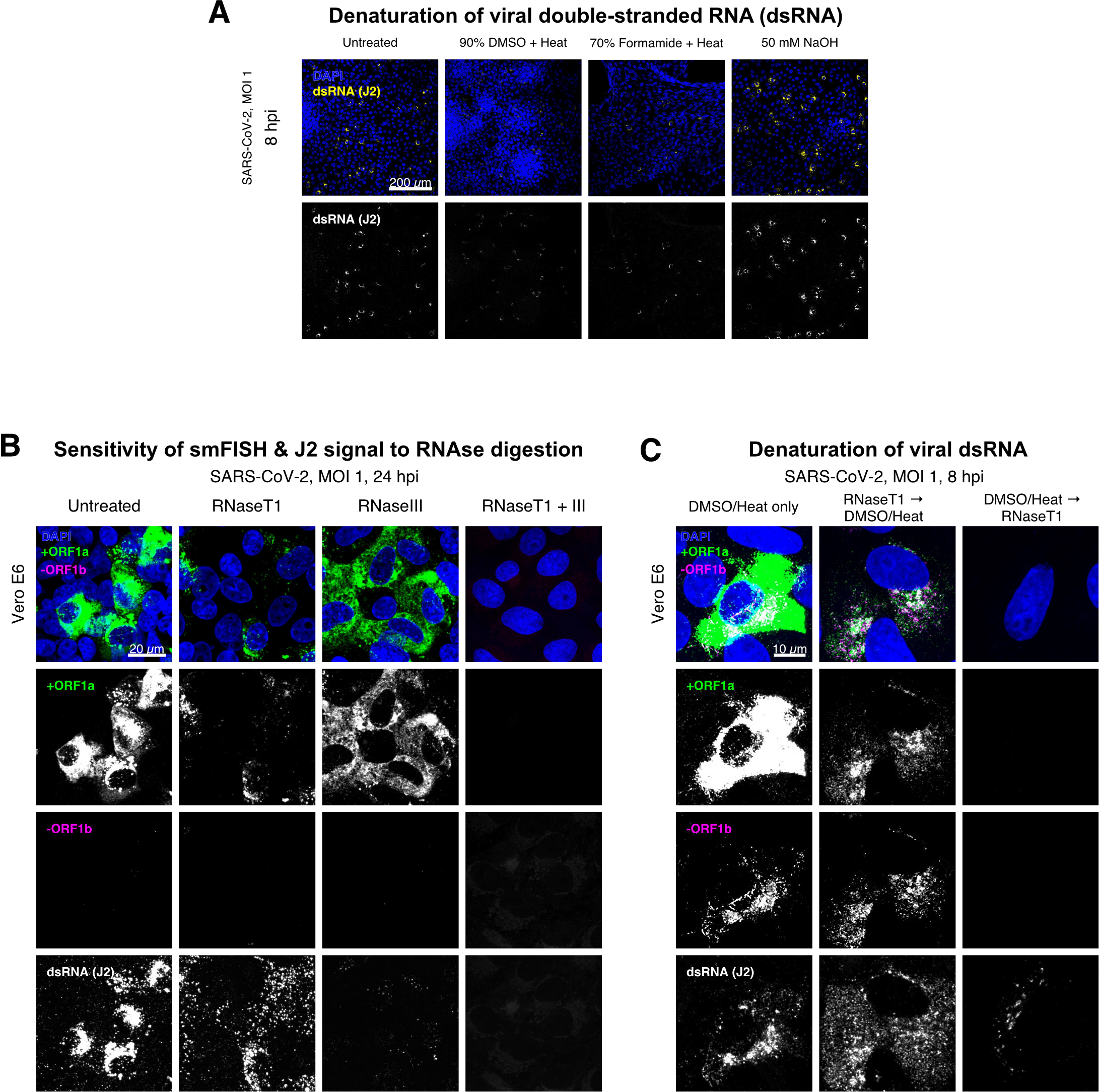
Denaturation of SARS-CoV-2 dsRNA for negative strand detection, related to Figure 2. (A) Comparison of double-stranded RNA (dsRNA) denaturation efficiency assessed by the reduction of anti-dsRNA (J2) immunofluorescence. Vero E6 cells infected with SARS-CoV-2 (VIC, MOI = 1) were fixed at 8 hpi and treated with DMSO, formamide or NaOH prior to immunostaining (see Materials and Methods). DMSO and formamide treatment was performed at 80°C. Representative low-magnification single slice confocal images are shown. Formamide treatment with heat resulted in cell detachment from coverslips. Scale bar = 200 µm. (B) Sensitivity of +ORF1a smFISH and J2 immunofluorescence signal to RNase digestion. Fixed infected cells (24 hpi) were treated with RNaseT1 and/or RNaseIII to digest single - stranded RNA and/or dsRNA, respectively, hybridised with +ORF1a probe and stained with J2. Representative full z-projected confocal images are shown, which are single-molecule contrast matched. Scale bar = 20 µm. (C) RNaseT1 digestion of denatured dsRNA. Fixed infected cells (8 hpi) were treated as follows: i) DMSO/heat only (left); ii) RNaseT1 then DMSO/heat (centre); or ii) in the reverse order of DMSO/heat and then RNaseT1 (right). Treated cells were hybridised with +ORF1a and -ORF1b probes (see Figure 2D for schematics) and stained with J2. RNaseT1 digestion followed by DMSO treatment shows that viral dsRNAs are resistant to RNaseT1 activity, but DMSO treatment preceding RNaseT1 suggests that the denatured dsRNA can be targeted by RNaseT1. Full z-projected confocal images are shown. Scale bar = 10 µm.

**Figure S3.**
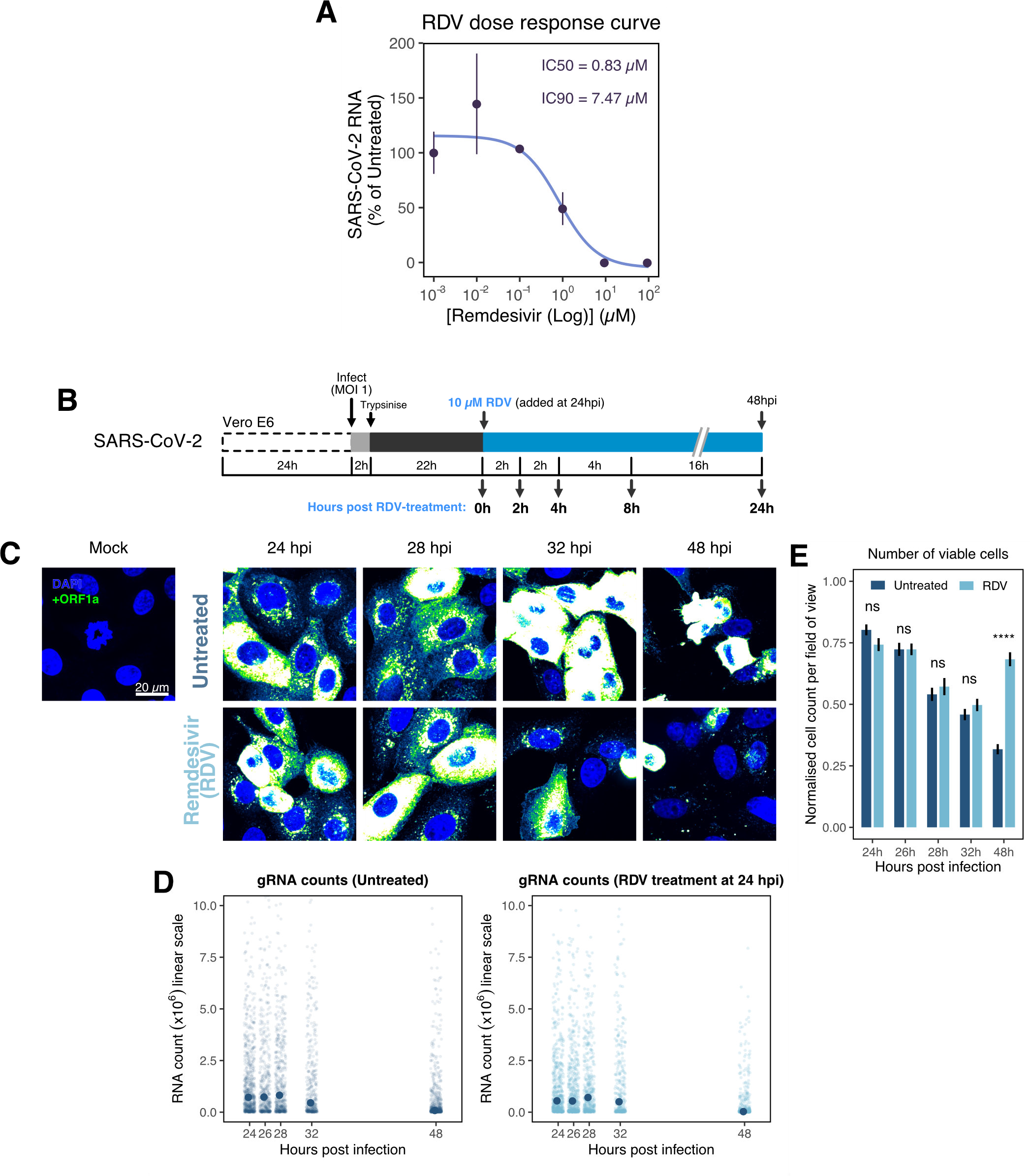
The dynamics and heterogeneity of SARS-CoV-2 RNA replication, related to Figure 3. (A) Dose response curve of remdesivir (RDV) versus SARS-CoV-2 RNA replication. Viral RNA was quantified using RT-qPCR in infected Vero E6 cells, targeting the ORF-N region. IC50 and IC90 values were estimated by fitting a nonlinear (weighted) least-squares model on the data (n=3). (B) Experimental design to profile SARS-CoV-2 replication kinetics in the late-stage infection. Vero E6 cells were seeded on cover-glass and, 24 h later, inoculated with SARS-CoV-2 (VIC strain, MOI = 1) for 2 h. Non-internalised viruses were removed by trypsin digestion and cells were fixed at timepoints shown for hybridisation with +ORF1a probe. In remdesivir (RDV) condition, the drug (10 µM) was added to cells at 24 hpi and maintained for the times shown. (C) Representative full z-projected confocal images of infected cells from the time series. Viral gRNA was visualised with +ORF1a probes. Images were contrasted to equivalent single molecule intensity. Scale bar = 20 µm. (D) Quantification of viral gRNA counts in untreated and RDV-treated cells. Reference single RNA molecule intensity was acquired using Bigfish in signal sparse region of the images. Viral gRNA counts were quantified by extrapolating single molecule intensity to 3D integrated intensity per cell. Data is represented as median ± S.E.M. (n=2). (E) The number of viable cells per field of view across the time series. Viable cells were quantified by counting non-condensed nuclei in randomly sampled field of views. Non-condensed nuclei counts were normalised to the average count values from uninfected ‘Mock’ condition. Student’s t-test. ****, p<0.0001. Data represented as mean ± S.E.M. (n=2).

**Figure S4.**
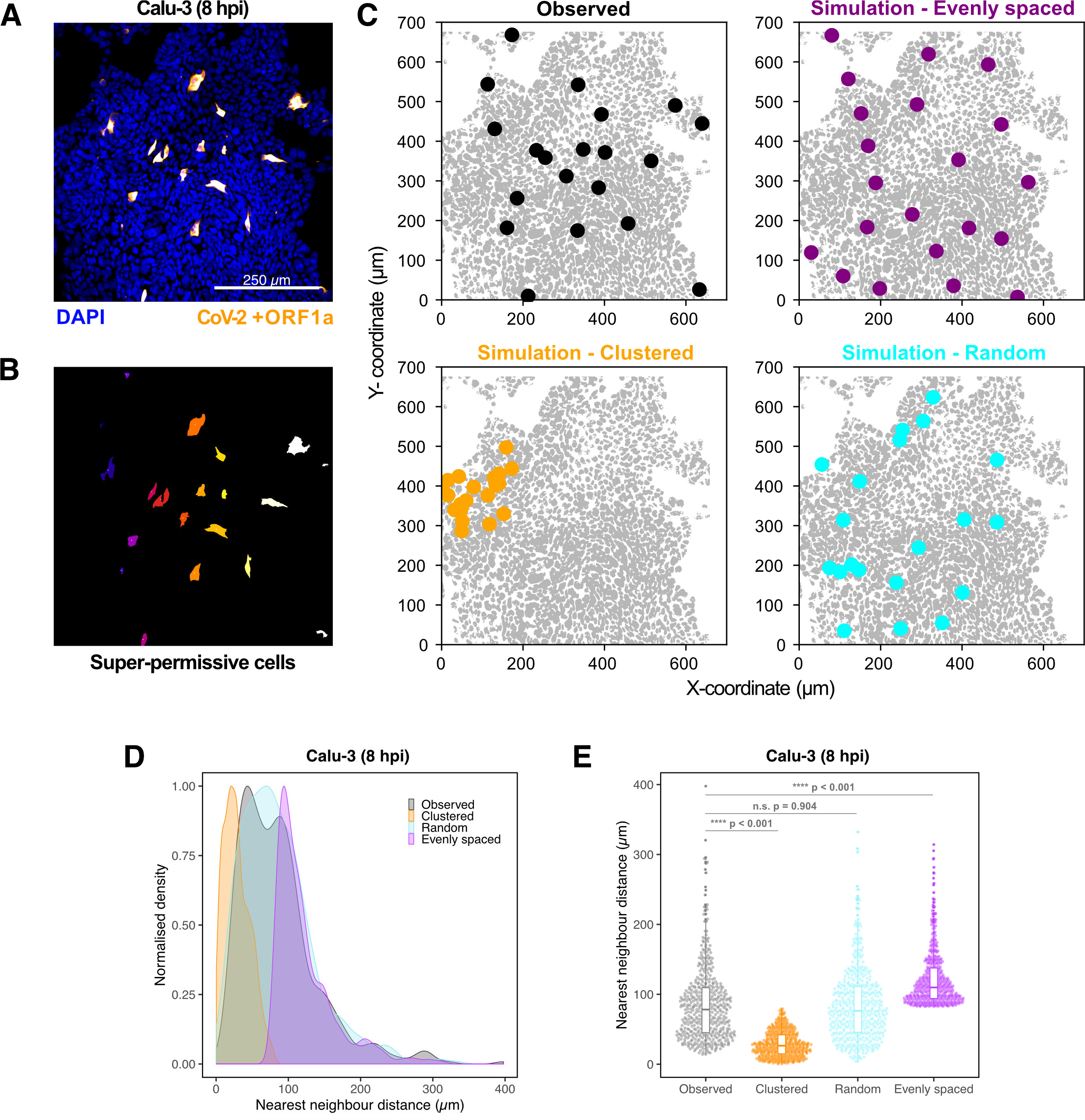
Spatial distribution of super-permissive SARS-CoV-2 infected cells, related to Figure 4. Spatial distribution analysis of ‘super-permissive’ SARS-CoV-2 (VIC) infected Calu-3 cells. Cells were infected with SARS-CoV-2 (VIC strain) at an MOI of 1, fixed at 8 hpi and hybridised with +ORF1a smFISH probe to visualise viral RNA. (A) Low magnification image (full z-projection) of infected Calu-3 cells showing a population of minority ‘super-permissive’ cells visible with smFISH. Scale bar = 250 µm. (B) 2D mask generated from spatial coordinates of super-permissive cells. (C) Observed distribution of super-permissive cells (*Observed*) and the example coordinates of the three modes of spatial distributions: i) evenly spaced, ii) clustered and iii) random. The simulations were confined to DAPI positive areas. (D) Density plot of nearest neighbour distances obtained from the spatial distribution simulation. Each mode of distribution was iterated 10 times per image. (E) Beeswarm plot of nearest neighbour distances obtained from the spatial distribution analysis. One-way ANOVA with post-hoc Tukey test. (n=3).

**Figure S5.**
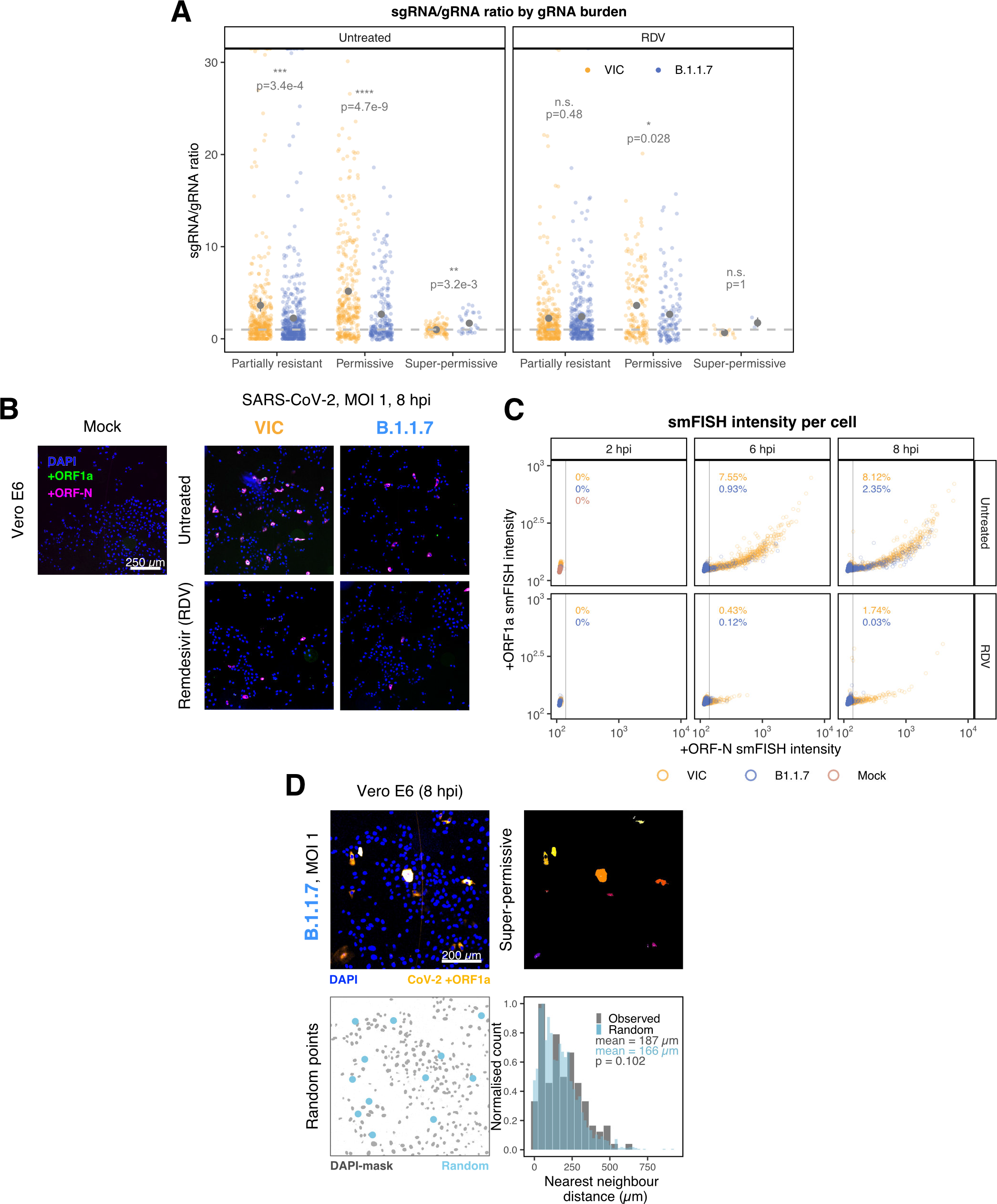
Delayed replication kinetics of B.1.1.7 SARS-CoV-2 variant, related to Figure 5. Vero E6 cells were seeded on cover-glass and 24 h later, inoculated with VIC or B.1.1.7 strain (MOI = 1) for 2 h. Non-internalised viruses were removed by trypsin digestion and cells were fixed at designated timepoints. In remdesivir (RDV) condition, the drug was added to cells at 10 µM during virus inoculation and maintained for the infection period. (A) Per cell ratio of sgRNA/gRNA counts grouped by gRNA burden classification as in (Figure 3D). smFISH quantification was performed as with (Figure 5H). Gray symbols represent mean ± S.E.M. The horizontal dashed line represents ratio of 1. (3). Mann-Whitney U test. n.s., not significant; *, p<0.05; **, p<0.01; ***, p<0.001; ***, p<0.0001. (B) Low-magnification (20x) z-projected (6 µm) images of +ORF1a & +ORF-N smFISH in infected Vero E6 cells. Scale bar = 250 µm. (C) Scatter plot showing high-throughput smFISH intensity quantification of +ORF1a and +ORF-N probes in VIC and B.1.1.7 infected cells. Each symbol represents a cell. Fluorescence density was measured from stitched 20x overview images, covering approximately ∼ 60% of the culture well area. At this magnification, smFISH fluorescence is only detectable in ‘super-permissive’ cells (>10^5^ vRNA). The percentage of super-permissive cells was calculated based on a gate which was set with +ORF-N signal using uninfected (Mock) condition signal as a threshold (vertical line) (n=3). (D) Spatial distribution analysis of B.1.1.7 infected Vero E6 cells showing ‘super-permissive’ phenotype. Low magnification overview image of ‘super-permissive’ cells visualised with +ORF1a probe (top left). 2D mask of super-permissive cells (top right). An example of randomly simulated points (bottom left). Same number of random points as ‘super-permissive’ cells were simulated 10 times per field of view. Histogram of nearest neighbour distances calculated between ‘super-permissive’ cells (Observed) and randomly simulated points (Random) (bottom right). (n=2).

**Figure S6.**
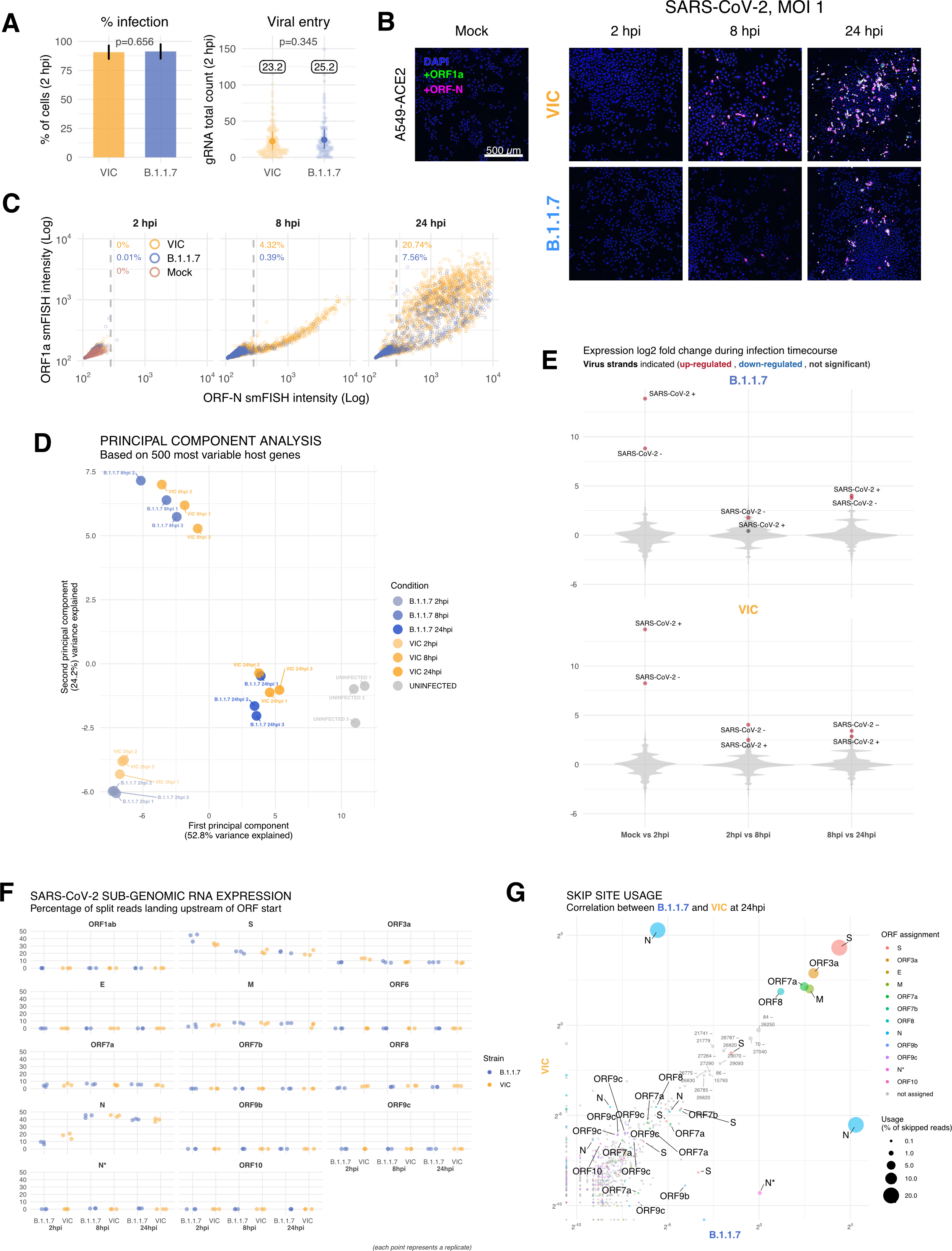
Transcriptomic landscapes of B.1.1.7 and Victoria and SARS-CoV-2 strains, related to Figure 6. A549-ACE2 cells were seeded and 24 h later, inoculated with VIC or B.1.1.7 strain (MOI = 1) for 2 h. Non-internalised viruses were removed by trypsin digestion and the cells were fixed at designated timepoints for smFISH or harvested for RNA-seq library preparation. (A) Comparing percentage of infected A549-ACE2 cells between VIC and B.1.1.7 at 2 hpi (left). Infected cells were determined by +ORF1a smFISH spot detection. Data represented as mean ± S.D. Comparison of viral gRNA counts at 2 hpi between the two strains (right). Each dot represents a cell. Data represented as mean ± S.E.M. and the labels represent average values. (B) Low-magnification z-projected (9 µm) images of SARS-CoV-2 infected A549-ACE2 cells. Cells were hybridised with +ORF1a & +ORF-N probes to visualise super-permissive cells. Scale bar = 500 µm. (C) Scatter plot showing high-throughput smFISH intensity quantification of +ORF1a & +ORF-N probes in infected cells. Each dot is a cell. Fluorescence signal density was measured from low-magnification overview image of entire culture wells. A gate was set with +ORF-N signal using uninfected (Mock) condition (dotted line) (2-4 hpi, n=2; 24 hpi, n=3). (D) First two components of a principal component analysis (PCA) performed on the 500 host genes showing the highest variance in RNA-seq. The infection timepoints (coloured) and control (gray) samples group into separate clusters but the samples the two strains remain close to one another. (n=3). (E) Violin plots showing fold-changes in the host transcriptome and viral RNA genome comparing consecutive timepoints separately for each of the two strains. Fold-changes for SARS-CoV-2 positive and negative strands are indicated as separate points and coloured according to the statistical significance of the change (red – upregulated relative to earlier timepoint, blue – downregulated relative to earlier timepoint, gray – no change). p-adjusted < 0.01, log2 fold change cutoff = 0. (n=3). (F) Expression of viral sgRNAs in each strain and different timepoints. Expression of each sub-genomic RNA is determined from split reads indicative of transcriptional skipping landing within 100nt upstream of annotated ORF start site, or until upstream ORF start codon if nearer. Percentage of all skip events is shown. (n=3). (G) Comparison of transcriptional skip site usage between the two virus strains at 24hpi. Assignment to viral genes is as in panel (F). Average across replicates is shown. (n=3).

