## Supplemental Data Item for "Absolute quantitation of individual SARS-CoV-2 RNA molecules: a new paradigm for infection dynamics and variant differences"

#### Supplementary Data Item

Multiple sequence alignment of SARS-CoV-2 variants of concern and variants of interest centred around transcriptional regulatory sequence motifs.

Occurrence of 10nt motif “CTAAACGAAC” in SARS-CoC-2 in relation to transcriptional skipping, ORF annotations, and mutations in different SARS-CoV-2 variants.

Top panels: Position of transcriptional skip ends (landing site) and their usage as percent of all skip events in B.1.1.7 (blue) and Victoria (orange) strains based on RNA-seq.

Middle panels: Multiple sequence alignment of SARS-CoV-2 reference strain, Victoria, and B.1.1.7 strains, as well as other variants of concern or variants of interest. 10nt motif “CTAAACGAAC” and its near variants are indicated by light blue boxes. Annotated ORF start and end codons are indicated in green and gray, respectively. Mutations relative to the reference strain are shown in yellow boxes. Bases with at least 0.1% skip site usage in B.1.1.7 (blue) or Victoria (orange) are indicated.

Bottom panels: Number of variants with mismatches relative to the reference strains at any given position.

#### 66 – 75 motif

Mutations, motifs, annotations, and skip site usage in Victoria and B.1.1.7

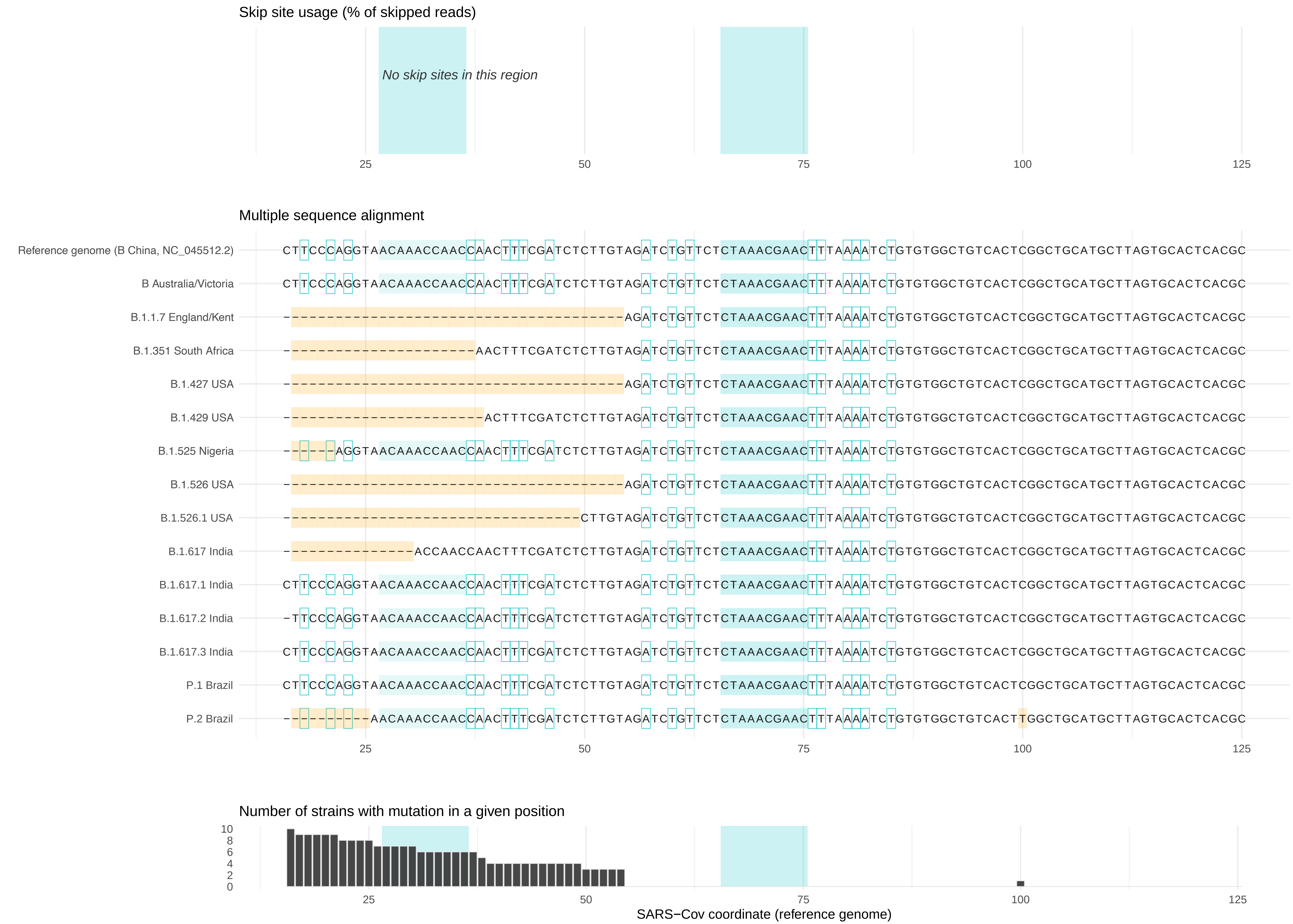


Highlighted regions indicate mutations

Annotations are based on reference genome

### 21552 – 21561 motif

Mutations, motifs, annotations, and skip site usage in Victoria and B.1.1.7

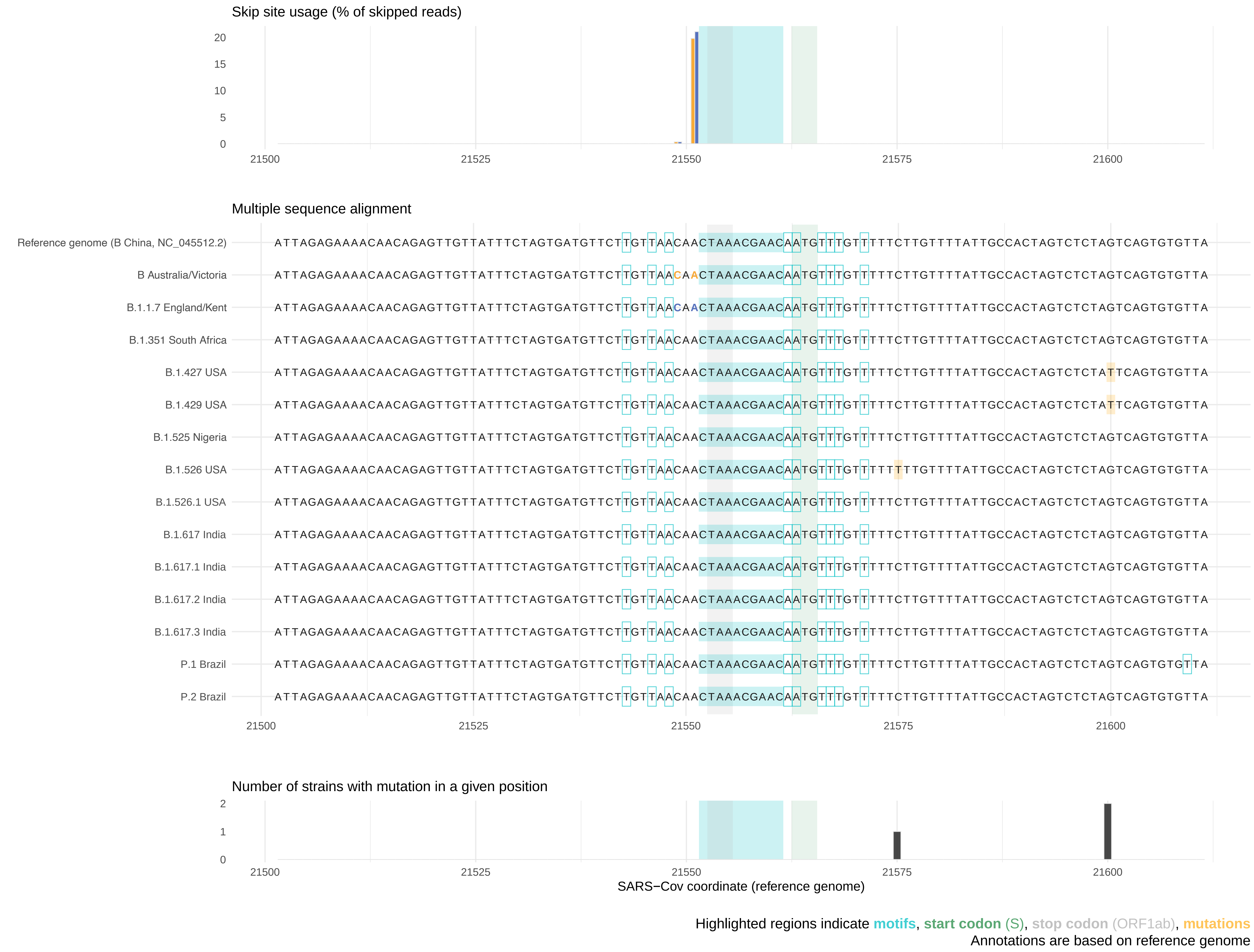

#### 25381 – 25390 motif

Mutations, motifs, annotations, and skip site usage in **Victoria** and **B.1.1.7**

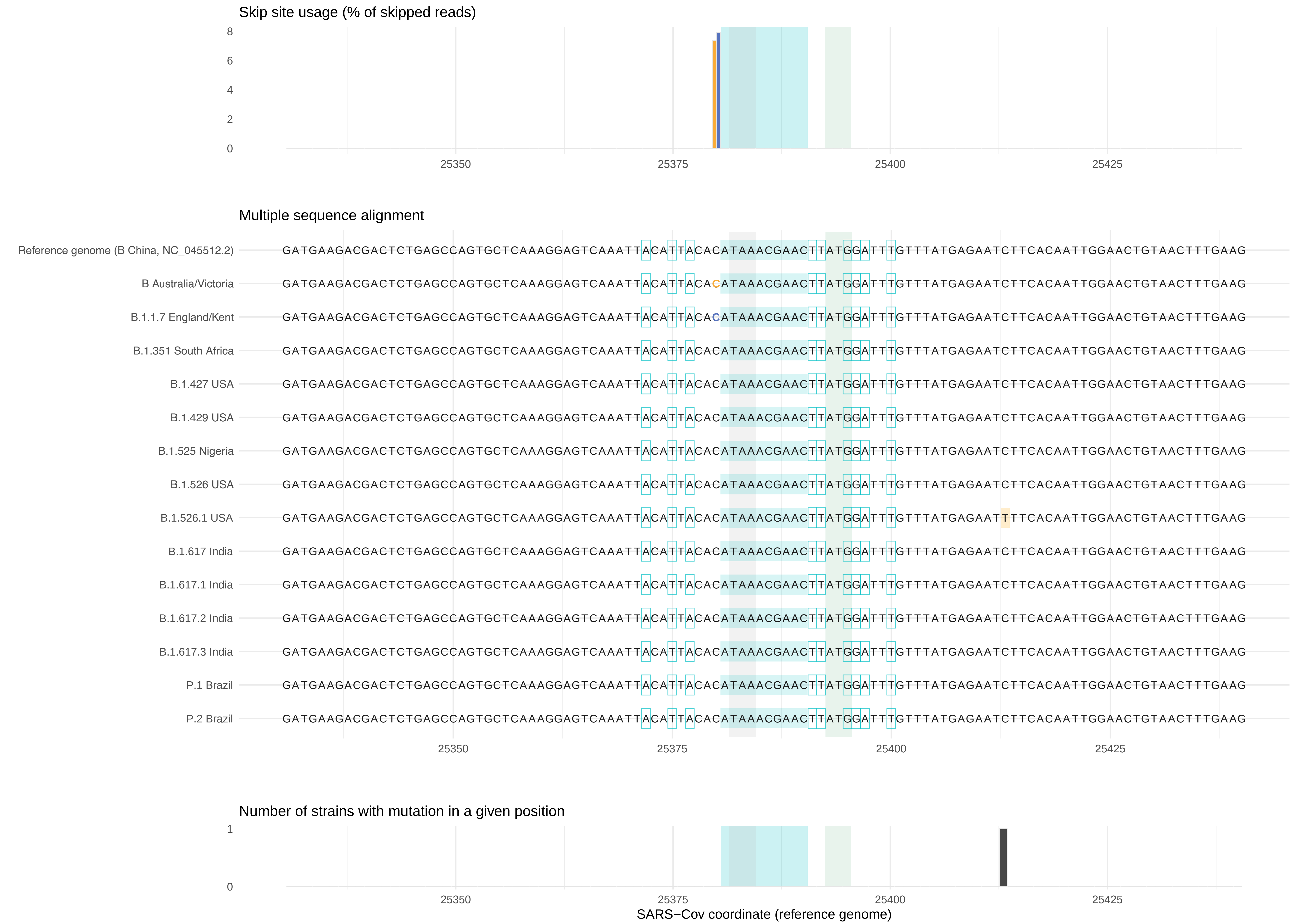


Highlighted regions indicate **motifs**, **start codon** (ORF3a), **stop codon** (S), **mutations**

Annotations are based on reference genome

#### 26469 – 26478 motif

Mutations, motifs, annotations, and skip site usage in Victoria and B.1.1.7

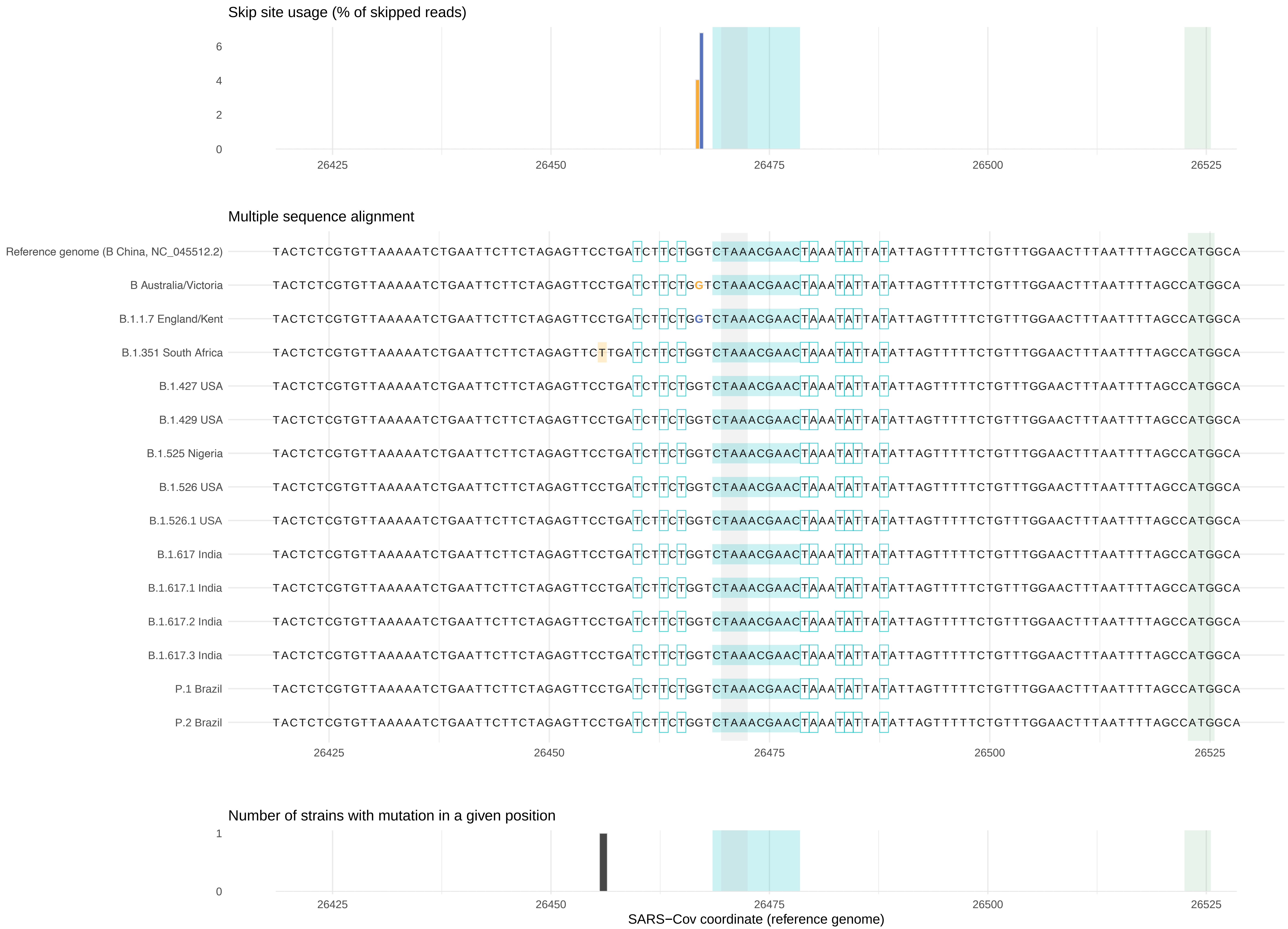


Highlighted regions indicate motifs, start codon (M), stop codon (E), mutations
Annotations are based on reference genome

### 27384 – 27393 motif

Mutations, motifs, annotations, and skip site usage in Victoria and B.1.1.7

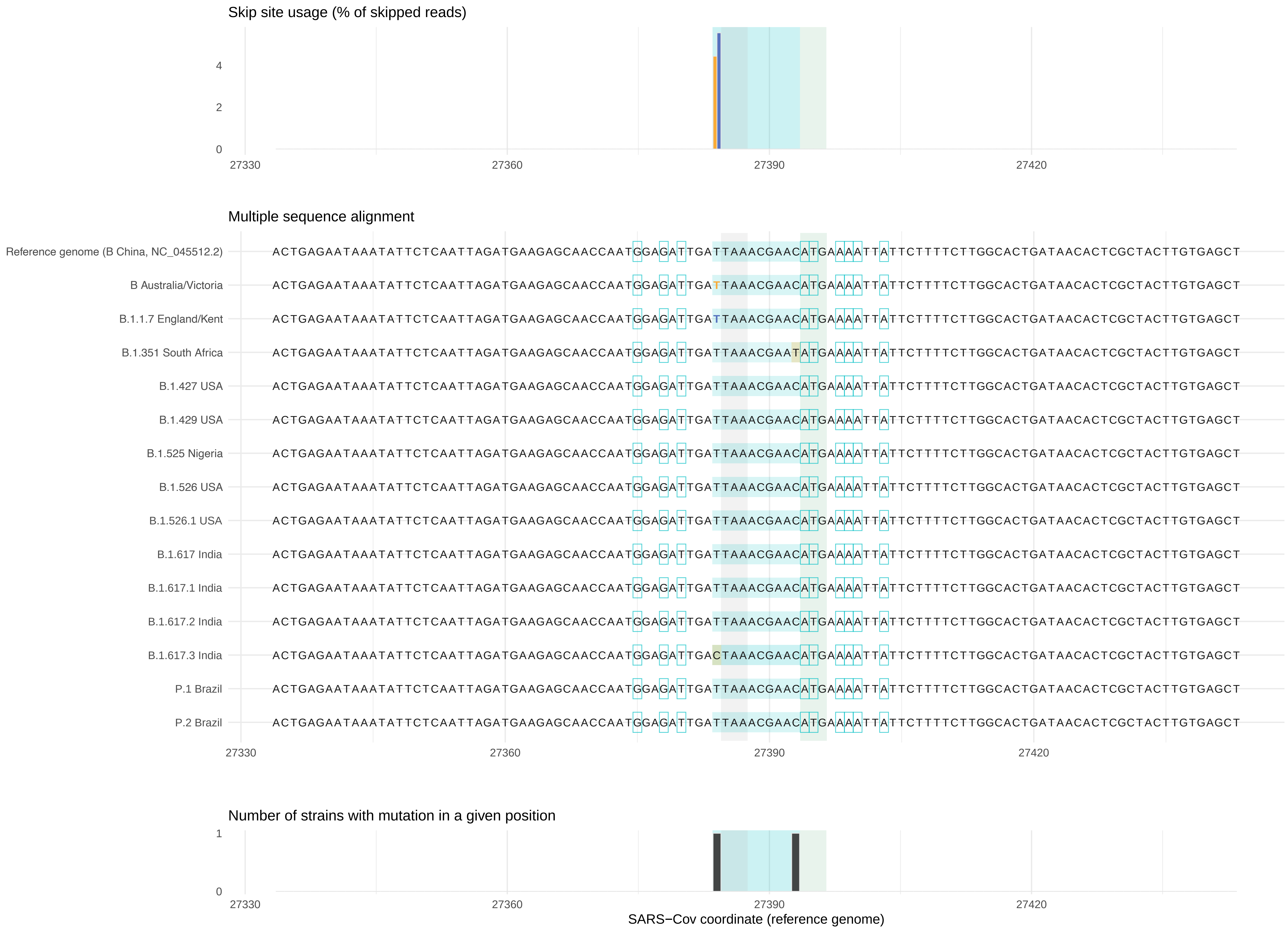


Highlighted regions indicate motifs, start codon (ORF7a), stop codon (ORF6), mutations
Annotations are based on reference genome

**27884 – 27893 motif**

Mutations, motifs, annotations, and skip site usage in Victoria and B.1.1.7

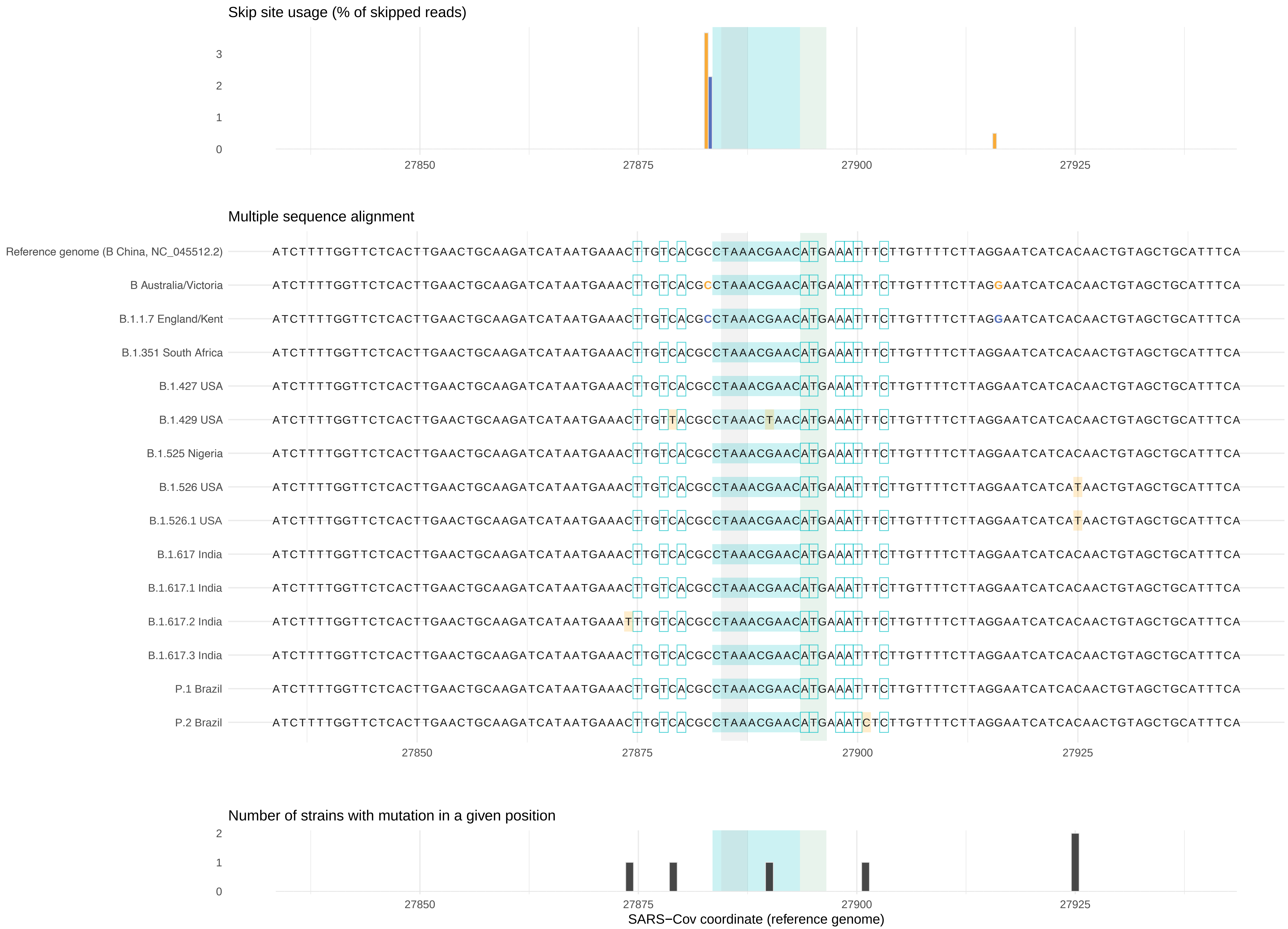


Highlighted regions indicate motifs, start codon (ORF8), stop codon (ORF7b), mutations

Annotations are based on reference genome

**28256 – 28265 motif**

Mutations, motifs, annotations, and skip site usage in Victoria and B.1.1.7

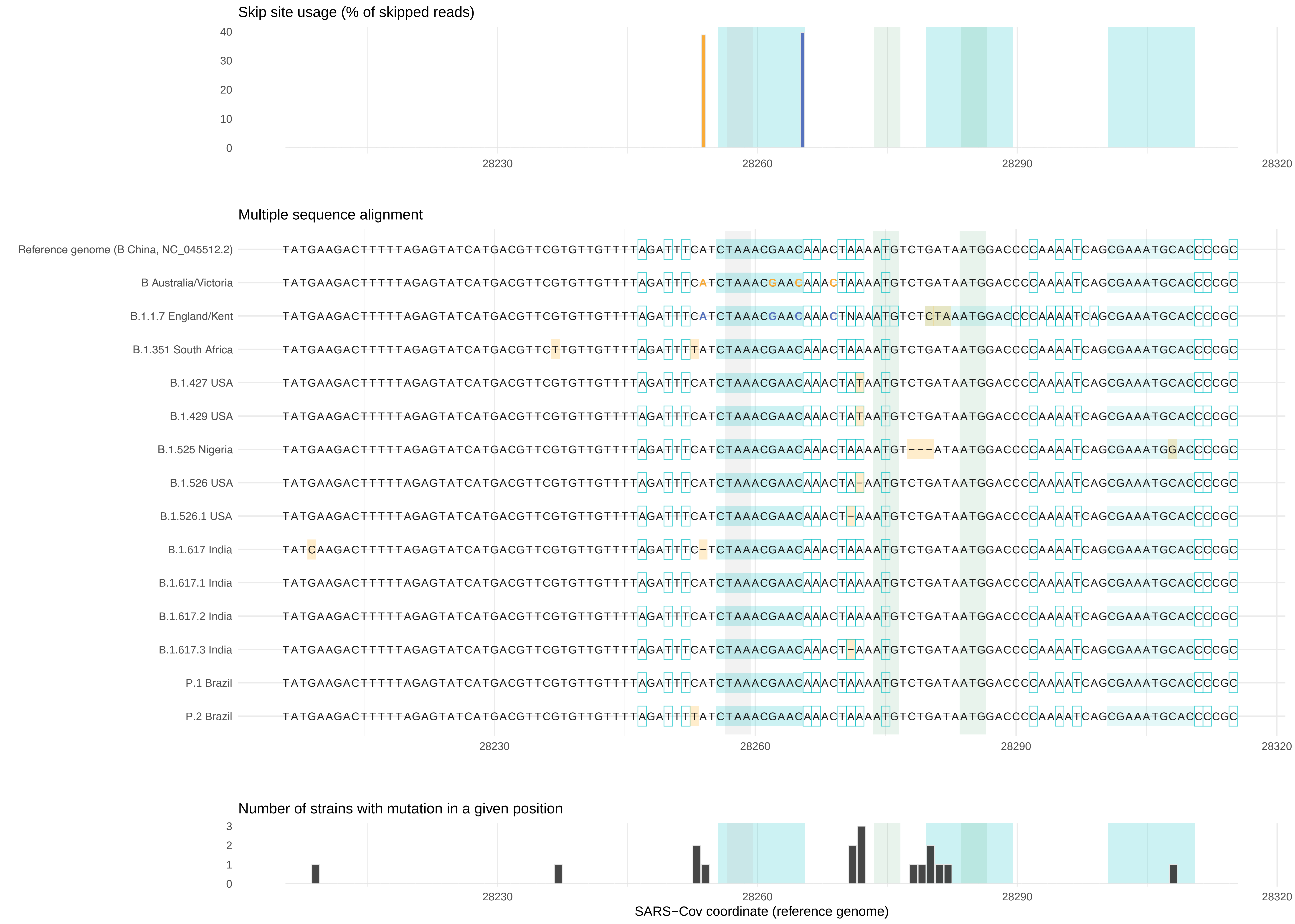


Highlighted regions indicate motifs, start codon (N, ORF9b), stop codon (ORF8), mutations

Annotations are based on reference genome

**28878 – 28887 motif**

Mutations, motifs, annotations, and skip site usage in Victoria and B.1.1.7

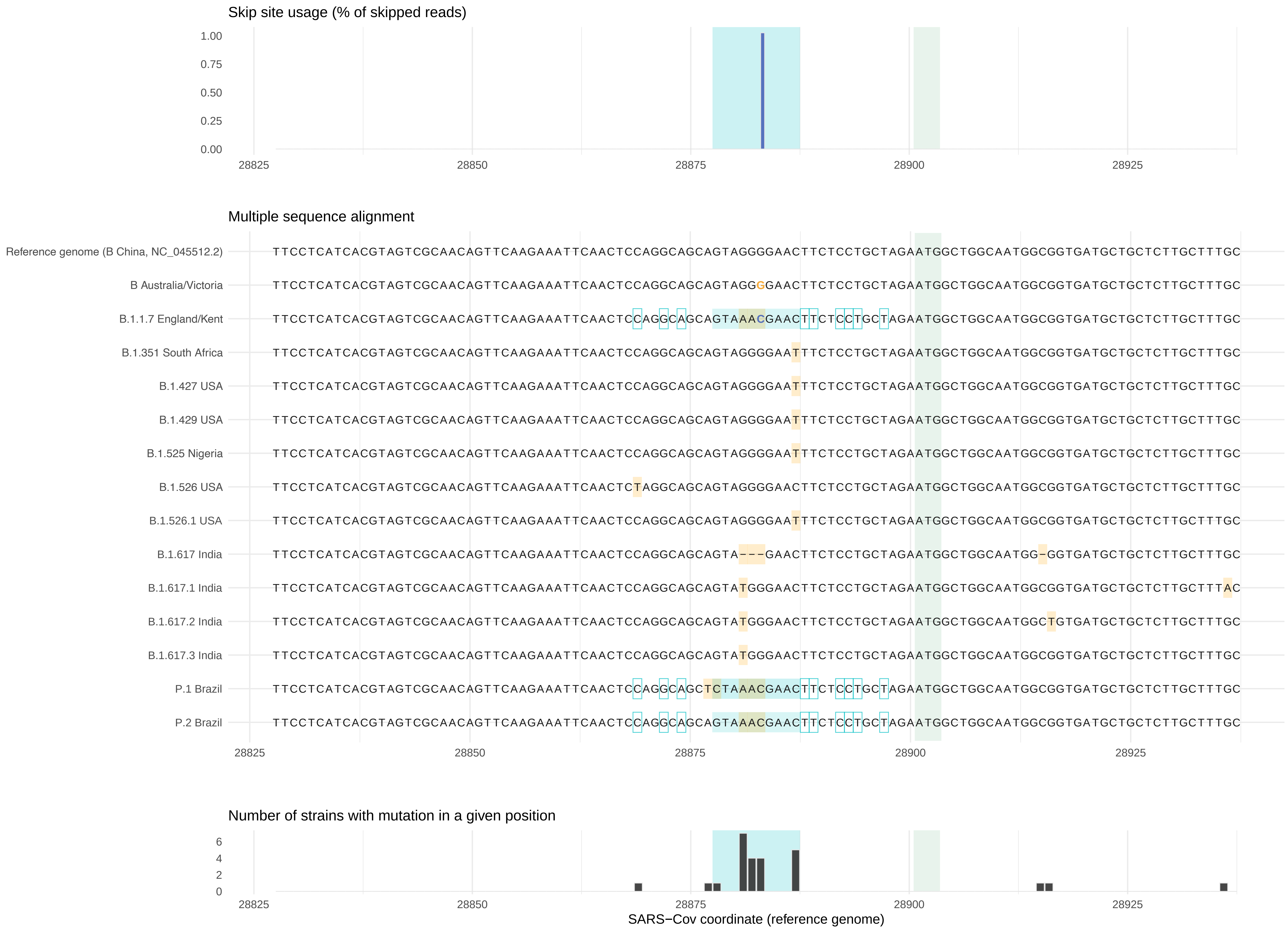


Highlighted regions indicate motifs, start codon (N*), mutations

Annotations are based on reference genome
